# Enhanced metabolic entanglement emerges during the evolution of an interkingdom microbial community

**DOI:** 10.1101/2024.03.30.587424

**Authors:** Giovanni Scarinci, Jan-Luca Ariens, Georgia Angelidou, Sebastian Schmidt, Timo Glatter, Nicole Paczia, Victor Sourjik

## Abstract

Metabolic interactions are common in microbial communities and are believed to be a key factor in the emergence of complex life forms. However, while different stages of mutualism can be observed in nature, the dynamics and mechanisms underlying the gradual erosion of independence of the initially autonomous organisms are not yet fully understood. In this study, we conducted the laboratory evolution of an engineered microbial community and were able to reproduce and molecularly track its stepwise progression towards enhanced partner entanglement. The evolution of the community both strengthened the existing metabolic interactions and led to the emergence of *de novo* interdependence between partners for nitrogen metabolism, which is a common feature of natural symbiotic interactions. Selection for enhanced metabolic entanglement repeatedly occurred indirectly, via pleiotropies and trade-offs within cellular regulatory networks. This indicates that indirect selection may be a common but overlooked mechanism that drives the evolution of mutualistic communities.

---

Microorganisms are typically part of communities that display a large taxonomic diversity and in which members are often linked through obligatory metabolite exchanges^1–4^. These interactions likely developed through a stepwise process, resulting in a gradual erosion of independence of the initially autonomous organisms^5,6^. Notably, similar processes likely guided eukaryogenesis^7,8^ and the emergence of obligate symbiotic interactions^5,6^. However, whilst communities composed of partners linked by various degrees of entanglement can be found in nature, the investigation of these evolutionary snapshots allows drawing only limited conclusions about the dynamics, molecular mechanisms and selection forces behind transitions towards increased cooperation^5,9^. Artificial synthetic communities may hence represent valuable models for the controlled observation of evolutionary processes in a relatively short time^10^. This approach was previously used to assemble mutualistic communities through passive metabolic interactions between microbial partners^11–17^, some of which could be evolved towards reinforced metabolite exchanges^18–24^. Nevertheless, the next phase of the community evolution predicted by ecological models^25–27^ - an increase in the interdependence mediated by the loss of traits - has not been experimentally reproduced so far, and there is still limited validation for theoretical frameworks proposed to explain how enhanced cooperation may be evolutionarily favoured over selfish behaviours^5,6,10,11,15,25,28,29^.

Since the most pronounced examples of metabolism reduction are observed for symbiotic interactions between prokaryotic and eukaryotic partners^6,30–33^, our study aimed to experimentally reproduce the transition towards increased cooperation by evolving an interkingdom <u>m</u>utualistic <u>co</u>nsortium between auxotrophs of *<u>E</u>scherichia coli* and *<u>S</u>accharomyces cerevisiae* (MESCo). We hypothesised that such interkingdom microbial community between a prokaryote and an eukaryote, where partners share no natural co-evolutionary history, might be more likely to undergo an evolutionary metabolic specialisation than previously studied purely bacterial^18,19,22^ or eukaryotic^20^ microbial communities.

## Experimental evolution of the MESCo communities leads to a rapid enhancement of growth

In order to identify a MESCo community that would be suitable for experimental evolution, we first tested the ability of different auxotrophs of *E. coli* and *S. cerevisiae* to complement each other for growth (**Extended Data Fig. 1a,b**). Despite previously reported challenges of their co-culturing^34^, we recently described conditions that enable stable propagation of a synthetic community between *S. cerevisiae* and *E. coli*^17^. Although growth was observed for multiple pairs of auxotrophs, the majority of tested communities turned out to be unsuitable for long-term evolution, either due to a community collapse or because of the spontaneous regaining of prototrophy by one of the partners. An exception was the MESCo community composed of *E. coli ΔhisG* and *S. cerevisiae Δarg1* strains, which could be stably propagated as a consortium in the selective <u>c</u>ross-feeding <u>m</u>inimal <u>m</u>edium (CF-MM) lacking histidine and arginine. Moreover, since *E. coli* is naturally able to co-aggregate with yeast via *type I* fimbriae^17^, we could investigate the possible influence of group selection^5,6,24,28^ enabled by such physical association between partners, by comparing the non-aggregating consortia containing fimbrialess (*ΔfimA*) *E. coli* (referred to simply as MESCo) with the aggregating consortia containing fimbriated (Fim^+^) *E. coli* (referred to as MESCo^Agg^).

Already after 15 regular transfers with a 1:10 dilution into fresh CF-MM, growth of the MESCo communities greatly improved compared to the ancestral (A) community (**Fig. 1a,b**). According to a total of approximately <u>50</u> generations of their passage under <u>c</u>ross-feeding conditions, these evolved communities were referred to as CF50. The aggregation status of the CF50 MESCo and MESCo^Agg^ (**Extended Data Fig. 1c**), as well as the auxotrophic status of both partners (**Extended Data Fig. 1d**), were retained throughout evolution. The growth of bacterial (*Ec*^CF50^) and yeast (*Sc*^CF50^) partners within the evolved communities was nearly proportional, and it was comparable between CF50 MESCo (**Fig. 1c**) and MESCo^Agg^ (**Fig. 1d**), despite initially negative effect of aggregation on growth of the ancestral MESCo^Agg^ community^17^. In our subsequent characterisation, we thus primarily focused on the detailed analysis of the MESCo communities evolved in the absence of aggregation.

**Fig. 1.**
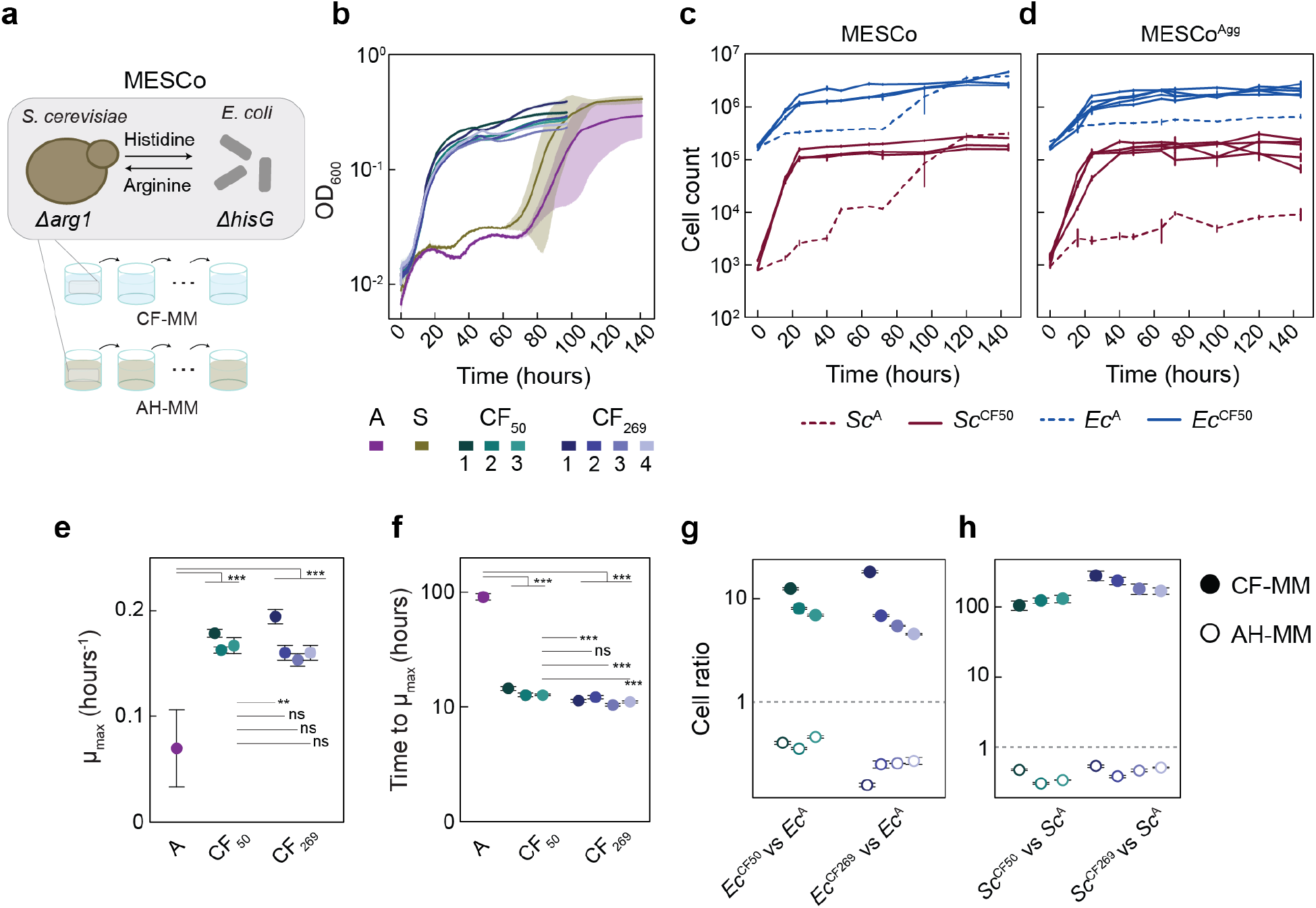
Design and experimental evolution of the MESCo communities (a) Schematic illustration of the design and experimental evolution of the bipartite MESCo communities between amino acid auxotrophs of *S. cerevisiae* and *E. coli*. Laboratory evolution was performed in the minimal media that were either selective for cross-feeding (CF-MM) or supplemented with arginine and histidine (AH-MM). (**b**) Growth of the non-aggregating ancestral community (labeled as A) or communities evolved either in CF-MM, for 50 (CF50) or 269 (CF269) generations, or in AH-MM (S), measured as optical density at 600 nm (OD600) in a plate reader. Colors and numbers denote different lines of evolution, and solid lines and shading indicate the mean values calculated from 3 biological replicates (*n*) and the corresponding standard deviation (SD). (**c,d**) Counts of *E. coli* (blue) and *S. cerevisiae* (red) cells in ancestral (dashed lines) or evolved (solid lines) communities, without (C) or with (D) aggregation, during growth in CF-MM. Cell counts were measured here and throughout using flow cytometry. Mean values of *n* = 3 biological replicates ± SD are shown. (**e,f**) Maximum growth rate (E) and time required to reach it (F) for growth curves in (B). Mean values of *n* = 3 biological replicates ± SD are shown. (**h,i**) Competition between evolved and ancestral community members, labelled with different fluorescent markers, which were inoculated in a co-culture at equal optical densities, and grown for 96 h either in CF-MM or in AH-MM as indicated. Cell ratio of the evolved bacterial (*Ec*^CF^; H) or yeast (*Sc*^CF^; I) lines, calculated as the ratio of the final cell counts for the indicated competing strains, normalized by their ratio at inoculation. A value of 1 represents absence of growth bias between the competing strains whilst values higher or lower than 1 indicate that the first strain respectively outcompetes or is outcompeted by the second one. Mean values of *n* = 5 biological replicates ± SD are shown. *p* values (ns = *p* > 0.05, ** = *p* < 0.01, *** = *p* < 0.001) reported in (e) and (f) are from a one-way ANOVA followed by Tukey post-hoc test.

The communities evolved under cross-feeding displayed both a moderate increase in the maximum growth rate (µmax) (**Fig. 1e**) and a dramatic reduction in the time that was required to reach µmax (**Fig. 1f**). In contrast, when both partners were co-evolved for approximately 100 generations in <u>m</u>inimal <u>m</u>edium containing both <u>a</u>rginine and <u>h</u>istidine (AH-MM), and thus without cross-feeding (referred to as S, for <u>s</u>upplemented), there was only a slight increase in µmax and no shortening of the time to reach µmax (**Fig. 1b** and **Extended Data Fig. 2a-c**). Thus, the reduction in the time to reach µmax appears to be specific for the co-evolution of community under cross-feeding. Consistently, there was a small but significant additional reduction in the time to reach µmax in three out of four communities obtained by subsequent evolution of CF50 3, up to a total of <u>269</u> generations (referred to as CF 269) (**Fig. 1f**), whereas µmax and the final cell densities only increased for one of the CF 269 communities (**Fig. 1b,e**). Notably, the ratio between both partners remained relatively stable over the course of evolution (**Extended Data Fig. 2d-g**).

When the ancestral and evolved communities were co-cultured in either CF-MM or AH-MM, both evolved *E. coli* (*Ec*^CF^) and *S. cerevisiae* (*Sc*^CF^) strongly outcompeted the ancestral strains (*Ec*^A^ and *Sc*^A^) in CF-MM, while being outcompeted in AH-MM (**Fig. 1g,h**). This confirms that the improved growth of the evolved MESCo communities in CF-MM is due to a specific advantage under conditions of cross-feeding, and this adaptation imposes a fitness cost in the absence of metabolic interdependency.

## A small set of mutations captures genetic changes in evolved communities

Sequencing the genomes of evolved populations of *E. coli* and *S. cerevisiae* revealed a small set of common mutations, appearing largely in the same sequential order in communities evolved without or with aggregation (**Fig. 2a** and **Supplementary Table 1,2**). The first mutations that became fixed in all evolved communities interrupted *argR*, which encodes the transcription factor that represses the biosynthesis and transport of arginine and histidine and regulates several other metabolic pathways in *E. coli*^35^ (**Fig. 2b**). Whilst inactivation of ArgR may enhance cross-feeding due to the increased production of arginine, such overproduction could also impose a metabolic burden. Indeed, the introduction of *ΔargR* (*Ec^ΔR^*) mutation into the ancestral *E. coli* strain led to a decreased growth rate (**Fig. 2c**), but higher levels of arginine (**Fig. 2d**) in the histidine-supplemented minimal medium. Given such negative effect on *E. coli* growth, strong positive selection on *argR* inactivation was thus counterintuitive. However, we hypothesised that these mutations may also derepress the *hisJQMP* operon encoding the histidine transporter^35^ (**Fig. 2b**), which was confirmed by the elevated activity of the *hisJ* promoter in *Ec^ΔR^*strain (**Fig. 2e**). Thus, the partner-serving inactivation of *argR* in *E. coli* is likely selected indirectly, due to an increased histidine uptake rather than enhanced cross-feeding. Consistent with positive selection on the histidine uptake, another set of *E. coli* mutations (in 4 out of 8 lines after 50 generations, and in all lines after 269 generations) directly affected the *hisJ* promoter region, resulting in a further increase in the promoter activity additionally to that provided by the *argR* deletion (**Fig. 2e** and **Extended Data Fig. 3a**).

**Fig. 2.**
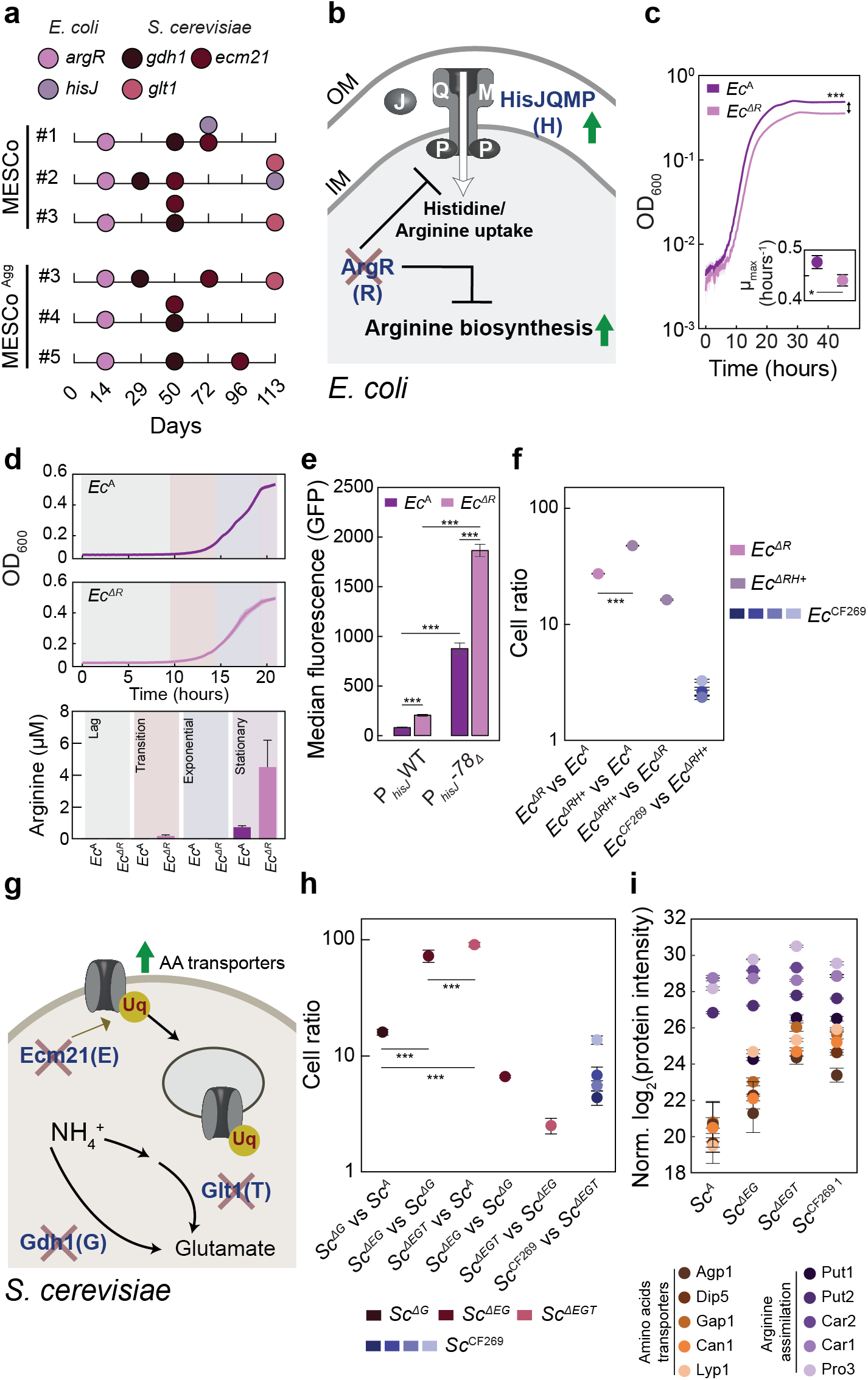
Common mutations fixed within the evolved MESCo communities (a) High-frequency mutations identified within genomes of populations of individual partners in communities evolved without (MESCo) or with (MESCo^Agg^) aggregation. Evolved lines are numbered throughout as in **Figure 1**. The identity and the time of first detection of the respective mutations within the evolving populations, assessed every two or three passages, are indicated. (**b**) Functions of affected proteins and expected impact of these mutations on *E. coli* partner (cross: nonsense mutations; up arrow: increased expression). (**c**) Growth of the ancestral *E. coli* (*Ec*^A^) and its *ΔargR* derivative (*Ec^ΔR^*) in CF-MM supplemented with histidine, with inset showing the maximum growth rate. Mean values of *n* = 5 biological replicates ± SD are shown. (**d**) Concentration of arginine measured in the supernatant at different growth phases from *Ec*^A^ or *Ec^ΔR^* cultures grown as in (c). Mean values of *n* = 3 biological replicates ± SD are shown. (**e**) Activity of the reporter plasmid, carrying either the wildtype (WT) *hisJ* promoter or one of its mutated versions (deletion in the nucleotide in position −78 from the start codon of *hisJ*; −*78Δ*) in front of the *gfp* gene, measured in either *Ec*^A^ or *Ec^ΔR^*background. Mean values of *n* = 3 biological replicates ± SD are shown. (**f**) Cell ratios of the indicated *E. coli* strains (*ΔRH+*: *ΔargR* and chromosomal *PhisJ* −*78Δ*) in pairwise competition in CF-MM, co-cultured with the ancestral yeast partner calculated as the ratio of the final cell counts for the indicated competing strains, normalised by their ratio at inoculation. Mean values of *n* = 4 (*Ec^ΔR^*, *Ec^ΔRH+^*) or 5 (*Ec* ^CF269^) biological replicates ± SD are shown. (**g**) Functions of affected proteins and possible impact of mutations on *S. cerevisiae* partner (cross: nonsense mutations). (**h**) Relative growth of the indicated *Sc* strains (A: ancestral; *ΔG*: *Δgdh1*; *ΔEG*: *Δecm21 Δgdh1*; *ΔEGT*: *Δecm21 Δgdh1 Δglt1*) in pairwise competition in CF-MM, co-cultured with *Ec^ΔR^* as a partner. Mean values of *n* = 5 to 6 biological replicates ± SD are shown. (**i**) Normalized protein intensity of indicated amino acid transporters regulated by *ecm21* and enzymes involved in arginine assimilation, in indicated yeast strains. Each strain was co-cultured in CF-MM for 36 h with *Ec^ΔR^* as a partner. Mean values of *n* = 4 biological replicates ± SD are shown. *p* values are reported in (**Supplementary Table 3**). *p* values (ns = *p* > 0.05, * = *p* < 0.05, ** = *p* < 0.01, *** = *p* < 0.001) reported in (C) and (F) are from a two-tailed *t*-test assuming unequal variance between the samples, in (H) and (K) are from a one-way ANOVA followed by Tukey post-hoc test.

The selective advantage of the identified mutations was confirmed by co-culturing different *E. coli* strains under cross-feeding conditions in the presence of the ancestral yeast strain (*Sc*^A^) (**Fig. 2f**). The ancestral *E. coli* strain (*Ec*^A^) was outcompeted by *Ec^ΔR^,* and even stronger by a strain carrying both the *ΔargR* and a mutation in the *hisJ* promoter (*Ec^ΔRH+^*), while *Ec^ΔR^* was outcompeted by the *Ec^ΔRH+^* strain. A similar result was obtained when all strains were co-cultured together with *Sc*^A^ under cross-feeding conditions, with the single mutants outcompeting the ancestral strain but being outcompeted by the double mutant strain (**Extended Data Fig. 3b**). Consistent with our mutation analysis, *Ec^ΔRH+^* strain largely recapitulated the phenotype of the evolved *E. coli*, being only slightly outcompeted by the *Ec*^CF269^ lines in the co-culture (**Fig. 2f**).

A similarly small set of common mutations was present at high frequencies in the evolved lines of *S. cerevisiae* (**Fig. 2a**), along with a number of low-frequency mutations (**Supplementary Table 2**). One prominent group of mutations introduced a premature stop codon in the gene *ecm21*, also known as *art2*. Ecm21 is a positive regulator of ubiquitination of several amino acid transporters, including the arginine transporter Can1, which promotes their endocytosis and subsequent degradation^36^ (**Fig. 2g**). Inactivation of Ecm21 likely benefits yeast under cross-feeding conditions because of the increased cell-surface levels of transporters and therefore increased uptake of amino acids, including arginine required by yeast. Notably, the emergence of similar mutations in *ecm21* was previously reported after the co-evolution between two different yeast auxotrophs^20^. Selection for the increased levels of Can1 may also explain the amplification of the entire chromosome V or of its region encoding the *can1* gene (**Extended Data Fig. 4**).

Another group of mutations in all evolved yeast lines interrupted *gdh1*, a gene encoding glutamate dehydrogenase. Gdh1 is the primary glutamate dehydrogenase used by *S. cerevisiae* growing on glucose^37^, and it catalyses one of the two major reactions for assimilation of ammonium (**Fig. 2g**). Although surprising given the presence of ammonium in the growth medium, the apparent selection for the loss of ammonium assimilation during evolution is further supported by the emergence of nonsense mutations (in 3 lines after 50 generations, and in all, except one, lines after 269 generations) in *glt1*, a gene that encodes the enzyme catalysing the second branch of direct ammonium assimilation (**Fig. 2a,g**).

These common mutations again conferred cumulative fitness advantage under cross-feeding conditions when introduced in the order of their appearance, evidenced by co-incubation of different yeast strains with *Ec^ΔR^* (**Fig. 2h**). This *E. coli* strain was chosen as a partner because *argR* mutations appeared in the community prior to any yeast mutations. Notably, there was a gradual decrease of the relative benefit for fitness provided by each subsequent mutation, with the deletion of *gdh1* giving the strongest benefit, followed by *ecm21* and then by *glt1*, which likely explains their order of fixation in the population. As for *E. coli*, these common mutations apparently capture most, but not all, of the beneficial changes in the evolved *S. cerevisiae* populations, since the *Sc*^CF269^ lines were moderately fitter than the triple knockout (*Sc^ΔEGT^*) strain (**Fig. 2h**).

The proteomics analysis confirmed that the evolved communities and the community formed by the *Sc^ΔEGT^* and *Ec^ΔRH+^* strains exhibit largely similar changes in protein levels compared to the ancestral community. These included upregulation of the HisJQMP transporter and proteins involved in the uptake and biosynthesis of arginine in *E. coli* (**Extended Data Fig. 5a** and **Supplementary Table 3**). Besides these common changes, both evolved lines, *Ec*^CF269 1^ and *Ec*^CF269 2^, showed a downregulation of the histidine biosynthetic pathway, as well as an upregulation of the outer membrane porin OmpF (**Extended Data Fig. 5b-d**) that is consistent with the mutations in the *ompF* promoter in these *Ec*^CF269^ lines (**Supplementary Table 1**). Similarly, changes in the *S. cerevisiae* proteome were largely overlapping between the evolved lines and the mutant community, including the expected upregulation of the arginine transporter Can1. However, the interpretation of these data was complicated by the difference in growth between the ancestral and evolved or mutant communities. We therefore analysed the proteome of different *S. cerevisiae* strains co-cultured with the same *Ec^ΔR^* partner, to ensure similar growth of all tested communities. These results confirmed elevated levels of Can1 in the *Sc*^CF269 1^ line and in the strains carrying key mutations, and further demonstrated the upregulation of several other amino acid transporters as well as proteins involved in arginine assimilation and nitrogen metabolism (**Extended Data Fig. 6a,b**).

In order to better understand the sequence in which mutations were fixed (**Fig. 2a**), we reconstructed communities between individual mutants. The inactivation of *argR* produced the most pronounced effect on growth (**Fig. 3a**), reflected in the strongly increased growth rate (**Fig. 3b**) and an even more dramatic reduction in the time to reach µmax (**Fig. 3c**). The effects of all subsequent mutations were less strong, with no or even negative impact on the growth rate or final density of the communities, but with a gradual reduction in the time to reach µmax. Given that a similar reduction was observed in the evolved communities (**Fig. 1b** and **f**), the time to reach µmax may be the main feature under evolutionary selection. The reconstituted communities carrying the major mutations were further able to phenotypically mimic the evolved lines (**Extended Data Fig. 7a-f**), including the ability to outcompete the ancestral community members under cross-feeding conditions, whilst being outcompeted in the supplemented medium (**Fig. 3d** and **Extended Data Fig. 7g,h**).

**Fig. 3.**
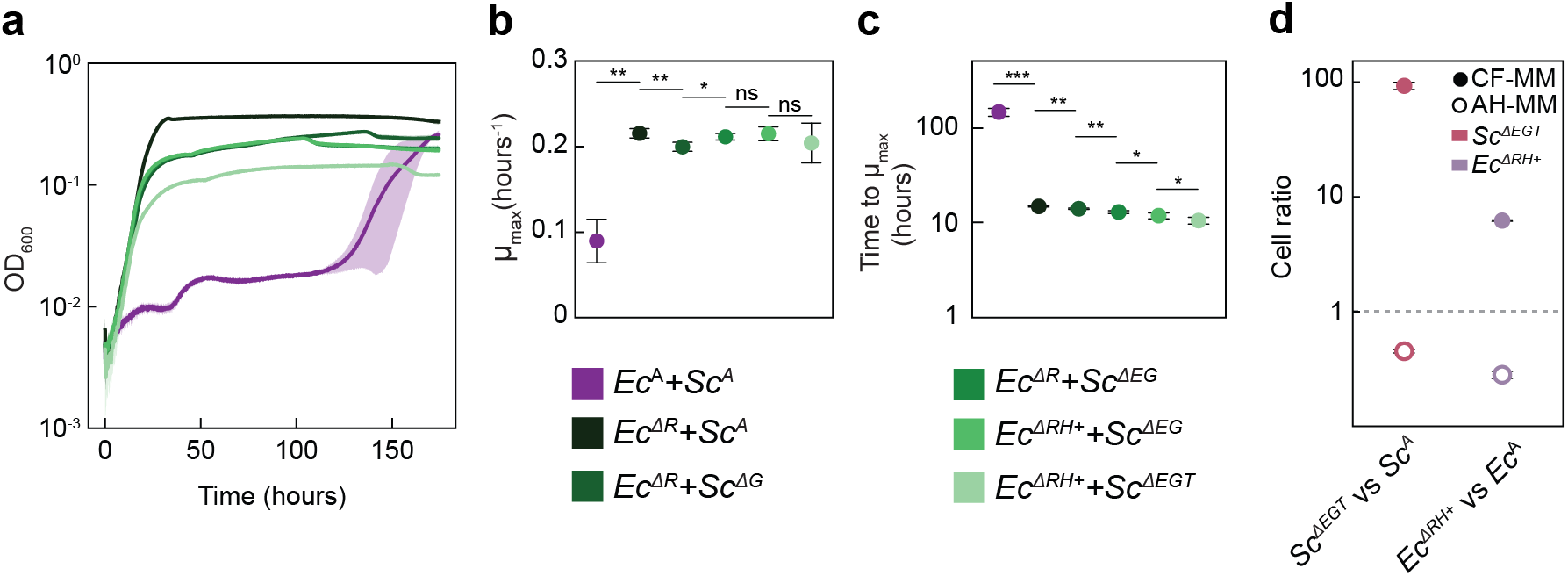
Impact of mutations on growth of MESCo communities. (**a-c**) Growth in CF-MM of co-cultures between indicated mutant strains representing consecutive stages of the community evolution (a), and corresponding maximal growth rate (b) and time to reach it (c). Mean values of *n* = 4 biological replicates ± SD are shown. (**d**) Cell ratio of yeast and bacterial mutants carrying the main mutations observed during evolution, in direct competition either in CF-MM or AH-MM with the ancestral counterparts. Mean values of *n* = 5 biological replicates ± SD are shown. *p* values (ns = *p* > 0.05, * = *p* < 0.05, ** = *p* < 0.01, *** = *p* < 0.001) in (b,c) are from a one-tailed *t*-test assuming equal variances between samples.

We also compared the growth of communities containing only individual mutations. While the deletion of *argR* or *ecm21* improved the growth of the community, the deletion of *gdh1* and the mutation of the *hisJ* promoter led to a reduction or cessation of community growth (**Extended Data Fig. 7i**), likely explains why these latter mutations could only be fixed at subsequent stages of the community evolution.

## Evolved yeast strains have a strongly reduced ability to directly assimilate ammonium

Whereas mutations related to the uptake and/or biosynthesis of histidine and arginine could enhance the pre-existing metabolic interactions within the MESCo communities, the fixation of yeast mutations affecting ammonium assimilation was unexpected. Nevertheless, further mutations in genes related to ammonium assimilation were observed in *Sc*^CF269^ lines (**Fig. 4a** and **Supplementary Table 2**), which include mutations in *gdh3* (paralogue of *gdh1*), in *mep1* and *mep3* that encode ammonium transporters, and premature stop codons in *gln3* that encodes the global transcriptional regulator of the nitrogen metabolism. Truncation of Gln3 was reported to produce a constitutively active version of this regulator^38^, and upregulation of Gln3 targets (including Can1) was indeed observed in the *Sc*^CF269 2^ yeast line carrying such truncation, when compared to the *Sc*^CF269 1^ line that retained the intact form of *gln3* (**Extended Data Fig. 8a**). The reduced ability of the evolved lines, as well as the corresponding yeast knockout strains, to assimilate ammonium was confirmed by growing them in arginine-supplemented minimal medium, either with or without ammonium. While the ancestral yeast strain exhibited much better growth in the presence of ammonium, the benefit from ammonium assimilation was largely reduced for both the evolved and mutant strains (**Fig. 4b-d** and **Extended Data Fig. 8b,c**), and such reduction was only marginal for control strains evolved in AH-MM without cross-feeding (**Extended Data Fig. 8d,e**).

**Fig. 4.**
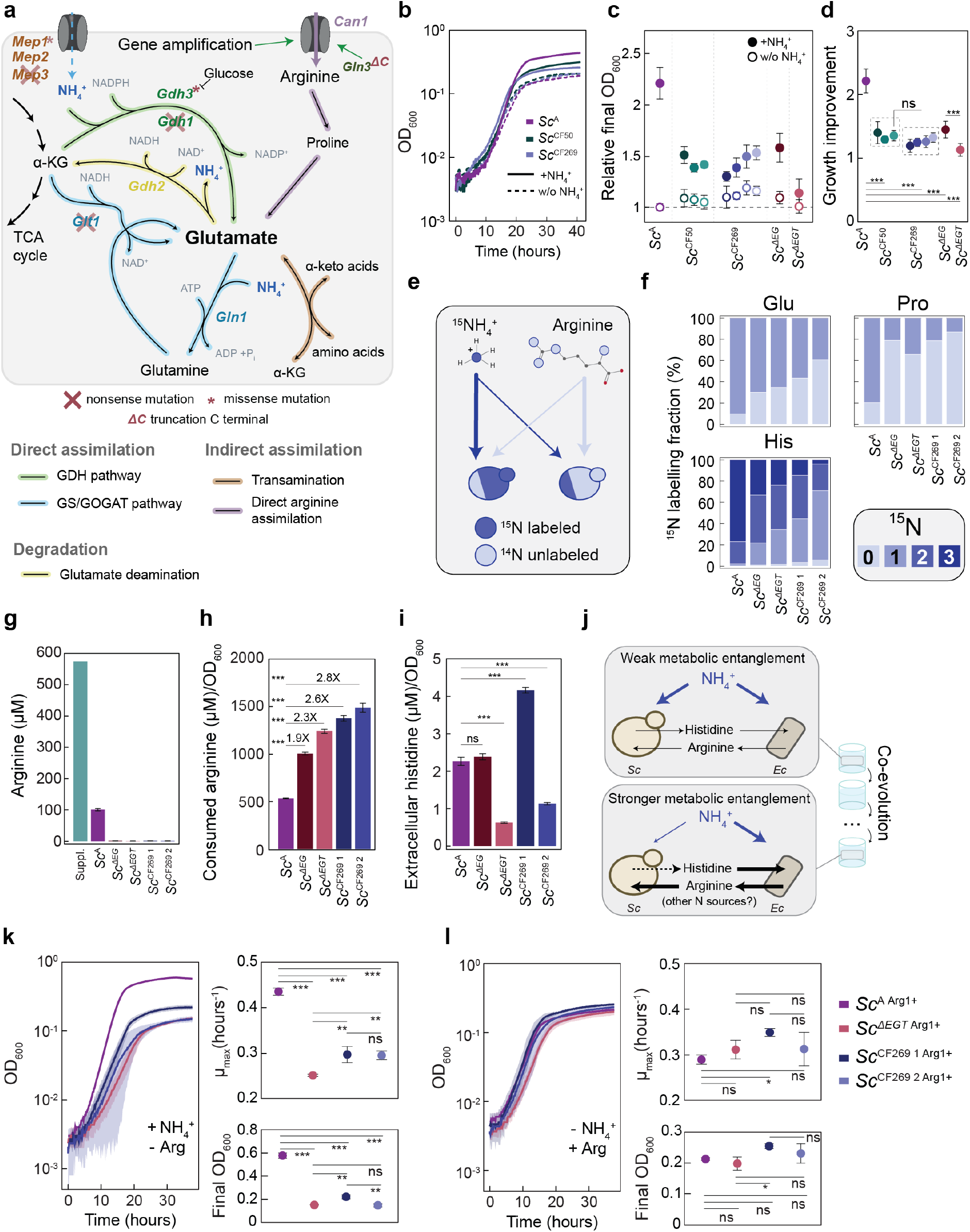
Modified nitrogen source preference of the evolved yeast partner (a) Schematic representation of the ammonium and arginine uptake and assimilation pathways in *S. cerevisiae,* highlighting corresponding mutations detected in the evolved yeast lines (cross: nonsense mutations; asterisk: missense mutations, ΔC: nonsense mutation causing truncation of Gln3). (**b-d**) Impact of ammonium on growth of the ancestral and evolved yeast lines in the arginine-supplemented minimal medium. Representative growth curves with or without addition of ammonium (b), final OD600 relative to the ancestral strain grown in the absence of ammonium (c), and growth increase due to the presence of ammonium (d) for indicated yeast strains. Mean values of (A, *n* = 11; *Sc^ΔEG^*, *Sc^ΔEGT^*, *n* = 9; *Sc*^CF50^, *n* = 5; *Sc*^CF269^, *n* = 5 to 8) biological replicates ± SD are shown. (**e**) Schematic illustration of the ^15^N labelling experiment. The ancestral (left) or mutant (right) yeast strains were grown in CF-MM supplemented with ^15^N-labelled ammonium and unlabeled (^14^N) arginine as nitrogen sources. Expected difference in the isotope labelling pattern is highlighted. (**f**) Average fraction of ^15^N-labelling detected in the indicated amino acids in stationary phase of the culture. Mean values of *n* = 4 biological replicates are shown, with SD (not shown) below 3% for all samples. (**g,h**) Concentration of arginine present in the supplemented CF-MM medium at inoculation (Suppl) and in the stationary-phase spent media of indicated yeast strains (g), and calculated consumption of arginine per unit of OD600 (h). Mean values of *n* = 4 biological replicates ± SD are shown. (**i**) Histidine levels per unit of OD600 detected in the spent media of indicated *S. cerevisiae* strains. Mean values of *n* = 4 biological replicates ± SD are shown. (**j**) Schematic representation of the evolved changes in the MESCo communities (see text for details). (**k,l**) Growth, maximal growth rate and final OD600 of indicated strains with restored arginine prototrophy in minimal media, with either ammonium (k) or arginine (l) as the primary nitrogen source. Mean values of *n* = 3 biological replicates ± SD are shown. *p* values (ns = *p* > 0.05, * = *p* < 0.05, ** = *p* < 0.01, *** = *p* < 0.001) reported in (d,h,I,k,l) are from a one-way ANOVA followed by Tukey post-hoc test.

When their preference for the source of nitrogen was directly tested, by growing yeast cells in the presence of the isotope-labelled ammonium and unlabeled arginine (**Fig. 4e**), the fraction of ^15^N-labelled proteinogenic amino acids, except arginine, was indeed much higher in the ancestral yeast strain compared to the evolved lines or to the mutant strains (**Fig. 4f** and **Extended Data Fig. 9a**). The effect was even more pronounced for the evolved lines compared to the *Sc^ΔEGT^*strain, likely because of the aforementioned additional mutations (**Fig. 4a** and **Supplementary Table 2**). The deletion of *gdh3* and *gdh2* in the *Sc^ΔEGT^* strain (labelled as *Sc^5KO^*) indeed resulted in a further reduction in the amino acid labelling (**Extended Data Fig. 9a**). Consistent with their increased reliance on arginine, the concentration of arginine in the supernatant of either evolved or mutant yeast strains was much lower compared to the ancestral yeast strain (**Fig. 4g**), implying an increase in consumption of arginine per OD600 unit (**Fig. 4h**).

We also tested the production of histidine, the metabolite provided by yeast to the bacterial partner within the consortium. However, in contrast to the general upregulation of arginine biosynthesis by the *E. coli* partner, only the *Sc*^CF269 1^ line, originating from the fastest-growing evolved community (**Fig. 1e**), showed increased abundance of histidine, and several other amino acids, in the supernatant (**Fig. 4i** and **Extended Data Fig. 9b**). The *Sc^ΔE^* mutant instead showed an unchanged level, and the *Sc^ΔEGT^* mutant and the *Sc*^CF269 2^ line showed even decreased levels of histidine, in contrast to the previous observation that *ecm21* mutations increased metabolite sharing in a yeast cross-feeding community^20^. Thus, the *Sc*^CF269 1^ line may have acquired additional mutation(s) that, possibly together with the *ecm21* mutation, enabled it to increase the cross-feeding of its partner.

We thus conclude that the evolution under cross-feeding conditions led to the increased reliance of yeast on arginine, supplied by its bacterial partner, as the primary nitrogen source instead of ammonium (**Fig. 4j**). This occurred despite the fact that ammonium was present in the medium during the entire course of evolution. To further verify this conclusion, we decoupled the use of arginine as the nitrogen source from the arginine auxotrophy in the yeast strains, by reintroducing the *arg1* gene to restore their arginine prototrophy. Although these “restored” strains could all grow in CF-MM without arginine, their growth on ammonium as the primary source of nitrogen was strongly reduced compared to the arginine-prototroph ancestral strain (**Fig. 4k**). In contrast, they grew equally well or even faster when arginine was provided instead of ammonium as the primary nitrogen source (**Fig. 4l**), or during residual growth on other supplements present in CF-MM (**Extended Data Fig. 10**).

## Restoration of prototrophy does not abolish the dependency of evolved yeast on the bacterial partner

Strains restored for the respective prototrophies further enabled us to directly test possible evolved dependencies within the community, beyond reliance on the exchange of arginine and histidine, by comparing growth of the prototroph strains in the absence and presence of the respective partner. Although cell counts of arginine prototrophs originating from the evolved yeast lines were lower compared to the restored ancestral strain (**Fig. 5a**), consistent with their reduced growth on ammonium (**Fig. 4k**), they grew significantly better in CF-MM in presence of *Ec^ΔRH+^* (**Fig. 5a,b**) (one-way ANOVA *p*=0.001, *R*^2^=0.48). Such enhancement was not observed for the restored ancestral strain or for the restored *Sc^ΔEGT^* prototroph, and it was weak for the restored 5KO prototroph, suggesting that *S. cerevisiae* indeed evolved additional dependence on the *E. coli* partner that may go beyond its increased reliance on arginine as the source of nitrogen. Supporting that, the final cell counts of the yeast prototroph strains did not differ significantly between communities containing either the ancestral or evolved *E. coli* (**Fig. 5c**), despite different levels of arginine excretion between these *E. coli* strains. Conversely, within the same community the evolved *E. coli* auxotroph partner does benefit more from the presence of the yeast prototroph (**Fig. 5d**), possibly due to its enhanced ability to scavenge histidine.

**Fig. 5.**
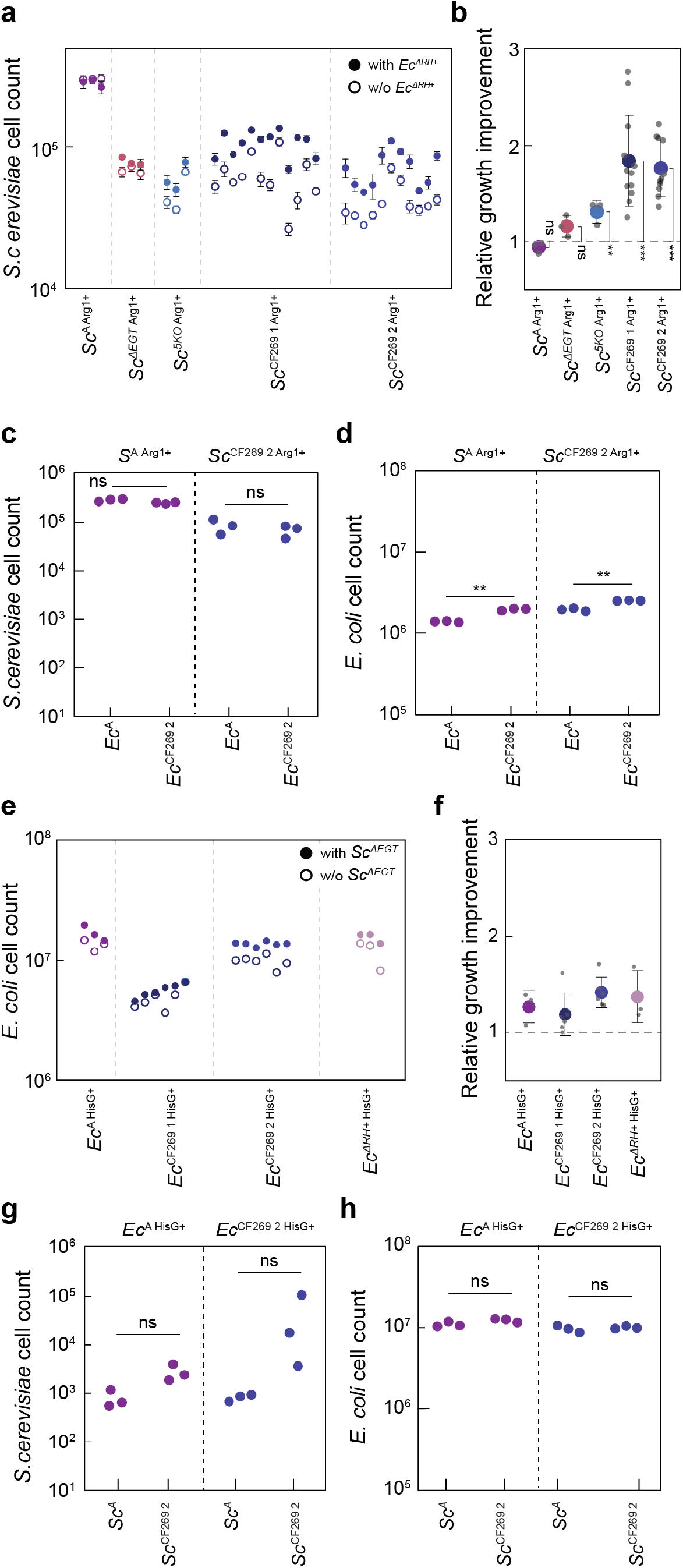
Partner dependencies of the evolved community members after restoration of their prototrophy. (**a.b**) Growth of indicated *S. cerevisiae* strains with restored arginine prototrophy in the presence or absence of *Ec^ΔRH+^* in CF-MM for 80 h. Mean values (*n* = 3) ± SD of the resulting cell counts (a), and the relative growth improvement in the presence of *E. coli*, calculated as the ratio between cell counts with and without bacterial partner (b) are shown. (**c,d**) Counts of yeast (c) and bacterial (d) cells in the co-cultures between indicated restored *S. cerevisiae* prototroph strains and either ancestral or evolved (*Ec*^269 2^) *E. coli* auxotrophs, grown as in (a). Each dot in all panels represents the cell count from individual transformants and is calculated as the average of two independent cultures. (**e,f**) The growth of indicated *E. coli* strains with restored histidine prototrophy in presence or absence of *Sc^ΔEGT^* in CF-MM for 80 h, with the resulting cell counts (e) and the relative growth improvement calculated as the ratio between cell counts with and without the yeast partner (f) are shown. Each dot represents the average of two biological replicates. (**g,h**) Counts of yeast (g) and bacterial (h) cells in the co-cultures between indicated restored *E. coli* prototroph strains and either ancestral or evolved (S*c*^269 2^) *S. cerevisiae* auxotrophs, grown in CF-MM for 80 h. Each dot represents the cell count from individual transformants and is calculated as the average from two biological replicates. *p* values (ns = *p* > 0.05, ** = *p* < 0.01, *** = *p* < 0.001) reported in (b) are from a one-sample *t*-test assessing for a difference from a value of 1 while in (c,d,g,h) are from a two-tailed *t*-test assuming unequal variance between the samples.

In contrast, when prototrophy was restored in *E. coli* strains, no significant difference in growth improvement was observed between the restored ancestral strain, the evolved lines and the *Ec^ΔRH+^*mutant (**Fig. 5e,f**) (one-way ANOVA *p*=0.28, R^2^=0.23), Furthermore, *E. coli* growth was not different when the partner yeast was either the ancestral or the evolved strain, and the evolved yeast did not benefit more strongly from the bacterial prototroph compared to the ancestral yeast strain (**Fig. 5g,h**). Thus, whereas yeast evolved additional dependencies on the bacterial partner beyond the originally engineered mutualism, *E. coli* enhanced its ability to profit from histidine but not from other metabolites provided by the yeast partner.

## Discussion

Ecological models predict that enhanced partner addiction should emerge from the co-evolution of interdependent organisms^27,39^, such as those exchanging essential metabolites in mutualistic and symbiotic communities^2–4,32^. In this study, we report experimental observation of an increase in metabolic entanglement during the experimental laboratory evolution of an engineered interkingdom mutualistic community between auxotrophs of *E. coli* and *S. cerevisiae* (MESCo). Natural symbiotic interactions often involve eukaryotic and prokaryotic organisms^6,32,33^, and compared to previously investigated monospecies consortia between either bacteria^13,18^ or yeast^14,20^, the two MESCo partners display larger differences in their metabolism and exometabolome profiles^40^. This is expected to favour cross-feeding interactions^41^ and potentially allows for a greater degree of mutual *de novo* adaptability.

Using this synthetic interkingdom community enabled us to mechanistically describe several characteristic steps in the progression of communities towards more efficient cooperation. This evolution firstly included the strengthening of pre-existing interactions, through the self-serving enhanced uptake of the exchanged metabolites by both partners, as previously observed in bacterial or yeast communities^16,20,21^. The cooperation was further promoted by the costly increase in sharing the partner-serving metabolite by *E. coli*, and at least in one instance also by yeast. Previous observations of the enhanced excretion of the partner-serving metabolites were primarily made in the context of laboratory evolution of spatially structured communities^18,19,24^, and the formation of multicellular clusters was even favoured by the co-evolution^23^. These instances of selection on cooperative traits could thus be interpreted as a consequence of local cooperation within small neighbourhoods^42^, favouring group selection that is normally assumed to be a prerequisite for the evolution of cooperation^5,6,10,11,15,28^.

In contrast, although group selection was specifically enabled in our experiments, the cooperative metabolite sharing rather emerged as a consequence of the pleiotropic mutations in the same regulatory component that simultaneously increased production of the partner-serving metabolite and uptake of the self-serving metabolite. While it has previously been demonstrated that pleiotropy can stabilise existing cooperation against the emergence of cheaters through regulatory coupling between cooperative and private traits^43–45^, the relevance of pleiotropy in the evolution of social traits has been questioned^46,29^. Nonetheless, in one case pleiotropy was suggested to explain the selection of the *ecm21* mutations in a mutualistic yeast community^20^, which may also apply to our experiments where the same yeast gene was mutated. Repeated instances of indirect selection during the short evolution of our MESCo communities suggest that pleiotropy may be a generally important, and previously underappreciated, factor in the evolution of sociality, promoting the emergence of social traits. Our results also indicate a mechanism that could favour selection of such pleiotropic over purely self-serving mutations, because of the observed negative impact of the latter on community growth.

Besides reinforcements of the pre-existing interactions, the evolved MESCo communities showed repeated emergence of a new level of dependency, with the yeast partner becoming increasingly reliant on *E. coli* for assimilation of ammonium, the primary nitrogen source in the medium during the co-culture evolution. This increased entanglement evolved through sequential inactivation of the major pathways of ammonium assimilation in yeast. Although the underlying selection pressure remains to be fully elucidated, the reduced ability of yeast to use ammonium may cause a rewiring of the nitrogen assimilatory pathways to enhance the uptake and consumption of arginine, thereby providing mutants with an increased scavenging ability for this metabolite, and thus with a competitive fitness advantage in a cross-feeding community. Additionally, the assimilation of ammonium under conditions of cross-feeding may cause some metabolic imbalance^47^, for example in the redox potential^48^, which could select for its inactivation. In either case, we conclude that this erosion of autonomy is again selected indirectly, as a consequence of regulatory trade-offs within the yeast metabolic network. Similar mechanisms may drive the emergent division of labour during the evolution of natural communities, as dependencies based on shared nitrogen-containing compounds are common in symbiotic interactions^32,49,50^.

## Methods

### Strain construction

*E. coli* knockout strains interrupted in different metabolic pathways were obtained from the Keio collection^51^ (**Supplementary Table 4**). Additional gene deletions in *E. coli* were introduced using pSIJ8 (**Supplementary Table 6**) as described^52^, after rescuing the strain from the kanamycin resistance (*kanR*) as described in^53^. Cassettes containing the *kanR* resistance cassette were amplified using respective gene deletion strains from the Keio collection, with flanking homology regions of 100 to 150 bp. For the introduction of the point mutations in the *hisJ* promoter region, cassettes containing the *neo-ccdB* were amplified from pKD45 using the primers GS_288 and GS_289 (**Supplementary Table 7**), which include a 50-bp homology from both sides for the desired region. This cassette was introduced in the desired *E. coli* strain as described^53^, and strains carrying this cassette were then transformed with a fragment containing the desired region amplified from the evolved lines, followed by selection on M9 minimal media plates containing rhamnose as sole carbon source. To distinguish strains during competition experiment using specific fluorescent markers, *E. coli* strains were transformed with pNB1 and pOB2 (for two-strain competition) or with pGS62-65 (for four-strain competition) (**Supplementary Table 6**).

*S. cerevisiae* strains were obtained from the respective knockout collection^54^ (**Supplementary Table 5**). Additional deletions in *S. cerevisiae* were obtained by transforming the desired strains with cassettes obtained by PCR amplification of the hygromycin B resistance from pH3FS (**Supplementary Table 6**), with flanking homology regions of 50 bp targeting the desired locus. Positive candidates were then transformed with the Cre-containing plasmid pPL5071 (**Supplementary Table 6**) and positive colonies were selected on minimal media plates lacking uracil and further screened via PCR to confirm the correct removal of the cassette containing the antibiotic resistance. Subsequently, after an overnight growth of positive candidates on complete minimal media at 30 °C, a selection on minimal media plates containing 5-fluoroorotic acid was performed to select for colonies lacking the pPL5071 plasmid. To distinguish strains during competition experiment via fluorescent markers, cassettes containing the *mNeonGreen* and *mTurquoise2* genes were amplified respectively from pMFM073 or pGS5 (**Supplementary Table 5**) and integrated in the *his3* locus of *S. cerevisiae*. For each competition experiment presented, the first strain listed is labelled with mTurquoise2 while the second always with mNeonGreen. For all the other experiments, strains expressing mTurquoise2 were used.

### Growth conditions

For *E. coli* pre-cultures, cells were inoculated directly from glycerol stocks into 5 ml lysogeny broth (LB), and if required the appropriate antibiotic was added. Pre-cultures were incubated at 37 °C for 16–18 h with shaking at 200 r.p.m. For *S. cerevisiae* pre-cultures, strains were firstly streaked from glycerol stocks on yeast extract peptone dextrose (YPD) plates, supplemented, when necessary, with the appropriate antibiotic. After incubation for 48 h at 30 °C, six colonies were inoculated in 5 mL YPD, supplemented, when necessary, with the appropriate antibiotic. Pre-cultures were incubated at 30 °C for 16–18 h with shaking at 200 r.p.m. Cells from 2 ml of the pre-cultures were collected by centrifugation, washed twice with phosphate saline buffer (PBS), suspended in 1 ml PBS and incubated for 5 h, either at 30 °C (*S. cerevisiae*) or 37 °C (*E. coli*). Cross-feeding experiments were performed in the low fluorescence (LoFlo) yeast nitrogen base (YNB) minimal medium (Formedium Ltd) buffered with 100 mM 2-(N-morpholino) ethanesulfonic acid (MES) (Roth) at pH 6.15, containing 2 % D-glucose as the carbon source and a mixture of 100 mg/l L-leucine, 20 mg/l L-methionine and 20 mg/l uracil to complement the auxotrophies present in the *S. cerevisiae* strains in the knockout collection^54^ (cross-feeding medium; CF-MM). Unless otherwise stated, both organisms were inoculated at OD600 of 0.05 each, values referring to a 1-cm cuvette. For competition experiment, each strain tested for each organism was inoculated in equal amount to a final total initial OD600 of 0.05. When supplements were used, a concentration of 100 mg/l of arginine and/or 20 mg/l of histidine were added to the minimal medium (supplemented minimal medium; AH-MM). Growth was measured as OD600 in a plate reader (Infinite 200 Pro, Tecan), by inoculating either the monocultures or the co-cultures in 48-well plates in 300 µl of the desired media and incubating the cultures for indicated time at 30 °C with 200 r.p.m. shaking. Growth plots were generated with a custom-written Python script and quantification of growth parameters was done using QurvE^55^ (non-parametric model). Statistical analyses were performed with GraphPad Prism v9.0.2.

### Experimental evolution

For the evolutionary experiments, co-cultures were inoculated in 1 ml of either CF-MM or AH-MM supplemented with 50 µg/ml kanamycin in 24-well plates and grown at 30 °C with 200 r.p.m shaking. For the evolution in CF-MM, the co-cultures were transferred into fresh medium at a ratio 1:10 every seven days for the first fifteen transfers (approximately 50 generations), at a ratio of 1:10 every 3.5 days between the 16^th^ and the 35^th^ transfer, and at a ratio of 1:200 until the 55^th^ transfer (for a total of approximately 269 generations). For lines evolved in AH-MM, co-cultures were transferred fourteen times at a ratio of 1:100 every 24 h (approximately 100 generations). In order to isolate different organisms from the co-cultures, communities were streaked respectively on YPD supplemented with 50 µg/ml of streptomycin to isolate yeast, and on LB supplemented with 50 µg/ml of nystatin to select for *E. coli.* The majority of colonies present on the plates were then pooled and grown in liquid cultures in the respective selective rich media, and these cultures were used to prepare glycerol stocks.

### Aggregation assay and microscopy

The aggregation assay was performed as described before^17^. Bacterial and yeast cells from pre-cultures, grown as described above, were washed twice with PBS and mixed together in 1 ml PBS in a 24-well plate (Greiner Bio-One GmbH) at a final OD600 of 0.7 for *S. cerevisiae* and 0.2 for *E. coli*. Plates were then incubated with shaking at 200 r.p.m. for 1 h at 30 °C and imaged using a Nikon SMZ745T stereo microscope.

### Sequencing

For Sanger sequencing, the genomic region of interest were firstly amplified by PCR (Q5-NEB), and the products were purified using the DNA Clean & Concentrator kit (Zymo Research). For the next-generation sequencing (NGS), genomic DNA extractions were performed using the NucleoSpin Microbial DNA Mini kit (Macherey-Nagel) following manufacturer’s instructions. In brief, pellets from 2 ml of LB-grown overnight cultures of *E. coli* were resuspended in 2 ml of the lysis buffer and homogenized (2 x 20 s at 6800 r.p.m. using Precellys Evolution, Bertin Technologies SAS). Pellets from 2 ml of YPD-grown overnight cultures of *S. cerevisiae* were resuspended in the lysis buffer, transferred to 400 µl suspension of HCl-treated glass beads (Merck KgAA) and vortexed with a Vortex Genie 2 (neoLab Migge GmbH) for 5 minutes at maximum speed. DNA concentration was quantified using a Qubit 4 Fluorometer (Thermo Fisher Scientific). For sequencing of CF50 communities, libraries were prepared using the Nextera XT DNA Library Preparation Kit (Illumina), and then sequenced using a Miniseq (Illumina). For CF269 communities, both library generation (NGS DNA Library Prep set-Novogene) and sequencing (Illumina NovaSeq 6000 S4 flowcell-Illumina) were performed by Novogene Co. Analysis of the sequencing data was performed using breseq^56^ and Integrative Genomics Viewer (IGV-version 2.8.9)^57^. Original fasta sequencing data are deposited in NCBI under the Bioproject PRJNA1049669 for CF50 and under the Bioproject PRJNA1051099 for CF269.

### Construction and analysis of promoter reporters

Plasmids carrying the *gfp* reporter under the control of different versions of *hisJ* promoter were constructed using the NEBuilder HiFi DNA Assembly (NEB). Primers used to amplify the backbone or the promoter regions from the evolved bacterial lines are reported in the primer list. *E. coli* strains transformed with the reporter plasmid were grown in 500 µl CF-MM supplemented with 20 mg/l histidine in a 48-well plate for 45 h at 30 °C, and afterwards fluorescence was measured via flow cytometry. In this case, the *Ec*^A^ rescued from *kanR* resistance was used as ancestral strain.

### Flow cytometry analysis

Flow cytometry measurements were performed using the BD LSR Fortessa SORP cell analyzer (BD Biosciences). A 488-nm laser line, with a power set to 20%, was used to determine both side scatter (SSC) and forward scatter (FSC) values, and combined with a 510/20 BP filter to detect GFP fluorescence. A 447-nm laser line combined with a 470/15BP filter was used to detect mTurquoise2 fluorescence, while the same laser line combined with a 586/15 BP was used to detect lss-mOrange. mCherry fluorescence was measured using a 561-nm laser line combined with 632/22 BP filter. *S. cerevisiae* and *E. coli* populations were distinguished by FSC and SSC, and the respective different strains used for competitions experiments were distinguished according to their respective fluorescent labelling (mCherry, GFP, mNeonGreen, mTurquoise2 and lss-mOrange for *E. coli,* mNeonGreen, mTurquoise2 for *S. cerevisiae*). Measurements were performed using the BD High Throughput Sampler (HTS) with a fixed flow rate set at 0.5 µl/s for an acquisition time of 20 s, with samples diluted in PBS to yield of 10^3^–10^4^ cell counts per second. If necessary, *S. cerevisiae*-*E. coli* aggregates were disrupted as described previously^17^, by diluting the community in PBS supplemented with 4% mannose followed by pipetting. The abundance of cells in the defined volume (10 µl) was inferred from the sample dilution, flow rate and sampling time. Flow cytometry data were analyzed using FlowJo (BD Biosciences).

### Proteomics sample preparation and liquid chromatography-mass spectrometry (LC-MS) measurements

To facilitate the collection and preparation of samples for the proteomic analysis, incubation was performed in trans-wells, where *S. cerevisiae* and *E. coli* partners are separated by a membrane (0.4 µm, Cellquart), which allows the metabolite exchange. Specifically, 3 ml of CF-MM containing *E. coli* partner at an initial OD600 of 0.083 were transferred into each well of a 6-well plate (SARSTEDT AG & Co. KG), a trans-well was inserted, and 2 ml of CF-MM containing *S. cerevisiae* partner at an initial OD600 of 0.125 was added. Cultures were grown at 30 °C with shaking at 110 r.p.m. Cells (equivalent to a total OD600 of 3.0) were harvested and washed three times with ice-cold PBS (15,000 g, 10 min, 4 °C) and resuspended in 300 μl of the lysis buffer containing 2% sodium lauroyl sarcosinate (SLS) and 100 mM ammonium bicarbonate. *E. coli* samples were then heated for 10 min at 90 °C, while *S. cerevisiae* samples were heated for 90 min at 90 °C. Samples were then ultra-sonicated for 10 seconds at maximum power (Vial Tweeter, Hielscher). Proteins were reduced with 5 mM tris (2-carboxyethyl) phosphine (Thermo Fisher Scientific) at 90 °C for 15 min and alkylated using 10 mM iodoacetamid (Sigma Aldrich) at 20 °C for 30 min in the dark. After centrifugation for 10 min at 13 000g, supernatants were transferred into a new tube. For *S. cerevisiae,* extracts were acetone-precipitated with a 4-fold excess of ice-cold acetone and incubation for 18 h at −20 °C, washed twice with methanol and dried for 10 min at room temperature. Dry pellets were then reconstituted in 200 μl lysis buffer. For both organisms, the amount of proteins was determined by bicinchoninic acid protein assay (Thermo Fisher Scientific).

For tryptic digestion, 50 µg of protein samples were incubated in 0.5% SLS and 1 µg of trypsin (SERVA Electrophoresis GmbH) at 30 °C overnight. After digestion, SLS was precipitated by adding a final concentration of 1.5% trifluoroacetic acid (TFA) (Thermo Fisher Scientific) followed by an incubation step of 10 min at room temperature. Peptides were desalted by using C18 solid phase extraction cartridges (Macherey-Nagel). Cartridges were prepared by adding acetonitrile (ACN), followed by equilibration with 0.1% TFA. Peptides were loaded on equilibrated cartridges, washed with 5% ACN and 0.1% TFA containing buffer and finally eluted with 50% ACN and 0.1% TFA.

Dried peptides were reconstituted in 0.1% trifluoroacetic acid and then analyzed using liquid-chromatography-mass spectrometry carried out on a Exploris 480 instrument connected to an Ultimate 3000 RSLC nano and a nanospray flex ion source (all Thermo Fisher Scientific). Peptide separation was performed on a reverse phase HPLC column (75 μm x 42 cm) packed in-house with C18 resin (2.4 μm; Dr. A. Maisch HPLC GmbH). The following separating gradient was used: 94% solvent A (0.15% formic acid) and 6% solvent B (99.85% acetonitrile, 0.15% formic acid) to 35% solvent B over 60 minutes at a flow rate of 300 nl/min.

MS raw data was acquired on an Exploris 480 (Thermo Fisher Scientific) in data independent acquisition mode with a method adopted from^58^. In short, Spray voltage was set to 2.3 kV, funnel RF level at 40, 275 °C heated capillary temperature, and 445.12003 m/z was used as internal calibrant. For DIA experiments full MS resolutions were set to 120.000 at m/z 200 and full MS, AGC (Automatic Gain Control) target was 300% with an IT of 50 ms. Mass range was set to 350–1400. AGC target value for fragment spectra was set at 3000%. 45 windows of 14 Da were used with an overlap of 1 Da. Resolution was set to 15,000 and IT to 22 ms. Stepped HCD collision energy of 25, 27.5, 30% was used. MS1 data was acquired in profile, MS2 DIA data in centroid mode.

Analysis of DIA data was performed using DIA-NN version 1.8^59^ using Uniprot databases for *Escherichia coli* or *Saccharomyces cerevisiae* to generate a data set specific spectral library for the DIA analysis. The neural-network based DIA-NN suite performed noise interference correction (mass correction, RT prediction and precursor/fragment co-elution correlation) and peptide precursor signal extraction of the DIA-NN raw data. The following parameters were used: full tryptic digest was allowed with two missed cleavage sites, and oxidized methionines and carbamidomethylated cysteins. Match between runs and remove likely interferences were enabled. The neural network classifier was set to the single-pass mode, and protein inference was based on genes. Quantification strategy was set to any LC (high accuracy). Cross-run normalization was set to RT-dependent. Library generation was set to smart profiling. DIA-NN outputs were further evaluated using the SafeQuant^60,61^ script modified to process DIA-NN outputs. The SafeQuant script was executed on the “report.tsv” file from DIA-NN analysis to sum precursor intensities to represent protein intensities. The peptide-to-protein assignment was done in SafeQuant with redundant peptide assignment following the Occam’s razor approach. Median protein intensity normalization was performed followed by imputation of missing values using a normal distribution function. Log-ratio and significance value (Student’s *t*-Test) calculation was performed as a basis for volcano plots with Perseus^62^. The mass spectrometry proteomics data have been deposited to the ProteomeXchange Consortium via the PRIDE^63^ partner repository with the dataset identifier PXD047443. Protein association network analysis and functional enrichment were performed with STRING^64^.

### Sample preparation for metabolite quantification

In order to quantify the arginine concentration in the *E. coli* supernatants, the cultures of the bacterial strains grown in 48 wells plates containing 300 µl of CF-MM supplemented with 20 mg/l histidine were filtered through a 15 mm 0.2 µm pore size reconstitute cellulose filters (Phenomenex Ltd) and flow-through samples were stored at −80 °C until measurement without any further treatment. In this case, the *Ec*^A^ rescued from the *kan* resistance gene was used as ancestral strain. For *S. cerevisiae* metabolites measurement in presence of ^15^N ammonium, cells were inoculated with an initial OD of 0.01 in 24-well plates containing 1500 ml of CF-MM without ammonium, supplemented with 100 mg/l arginine and 5g/l 98 % ^15^N (NH4)2SO4 (Merck KgAA), and grown at 30 °C until stationary phase. For metabolite quantification from supernatants, cultures were filtered and stored as above. For proteinogenic amino acid hydrolysis and extraction, samples were adjusted to equal biomass according to OD600, cells were collected by gentle centrifugation (3000 g, 10 min), and the pellets were washed three times with PBS. Washed pellets were suspended in 6N HCl solution and transferred to glass vials with conical base (ROTILABO-Carl Roth) and stored at 98 °C for 6 h. Samples were then dried under a nitrogen stream, suspended in 250 µl double distilled water and transferred into clean Eppendorf tubes. These were centrifuged at maximum speed for 10 minutes and the supernatants were transferred into clean Eppendorf tubes and stored at −80 °C until measurement without any further treatment.

### Metabolite quantification via LC-MS

Both quantitative and qualitative determination of the target metabolites were performed using HRES LC-MS. The chromatographic separation was performed on a Vanquish HPLC System (Thermo Fisher Scientific) using a ZicHILIC SeQuant column (150 × 2.1 mm, 3.5 μm particle size, 100 Å pore size) connected to a ZicHILIC guard column (20 × 2.1 mm, 5 μm particle size) (Merck KgAA), with a constant flow rate of 0.3 ml/min. The temperature was maintained at 25 °C. The two mobile phases were a solution of 0.1 % Formic acid in 99:1 water:acetonitrile (Honeywell research chemicals) as mobile phase A, and a solution of 0.1 % formic acid 99:1 acetonitrile:water (Honeywell research chemicals) as phase B. The injection volume used per each sample was set to 5 µl. The following steps and linear gradients were used for the mobile phase profile: 0 – 8 min from 80 to 60 % B; 8 – 10 min from 60 to 10 % B; 10 – 12 min constant at 10 % B; 12 – 12.1 min from 10 to 80 % B; 12.1 to 18 min constant at 80 % B. ID-X Orbitrap mass spectrometer (Thermo Fisher Scientific) was used in positive mode with a high-temperature electrospray ionization source and the following conditions: H-ESI spray voltage at 3500 V, sheath gas at 50 arbitrary units, auxiliary gas at 10 arbitrary units, sweep gas at 1 arbitrary units, ion transfer tube temperature at 350 °C, vaporizer temperature at 350 °C. Detection was performed in full scan mode using the orbitrap mass analyser at a mass resolution of 60 000 in the mass range 50 – 250 (m/z). Extracted ion chromatograms of the [M+H]+ forms were integrated using Tracefinder software (Thermo Fisher Scientific). For the reported intensity levels of the different amino acids, values were obtained by summing the area under the peaks from LC-MS measurements for the different isotopologues. Absolute concentrations were then calculated based on external calibration curves.

### Data and materials availability

Original proteomics and sequencing data have been deposited in public repositories as indicated in Materials and Methods. All the other data are available in the main text or in Supplementary Information. All materials are available from the corresponding author upon request. The proteomics data have been deposited to the ProteomeXchange Consortium via the PRIDE partner repository with the dataset identifier PXD047443 and are accessible for reviewing under username: and password: P75czZut.

## Acknowledgments

We thank Jörg Kahnt for the support with the proteomics analysis, Silvia Gonzalez Sierra for the support with the flow cytometry, and Elif Elçin and Paushali Chaudhury for the support with the NGS sequencing. We thank Julian Pietsch for insightful discussions. We thank John S. Parkinson for providing the materials and the protocol for gene replacement in *E. coli*. This research was funded by the Max-Planck-Gesellschaft.

## Author contributions

G.S. and V.S. conceived and designed the study. G.S., J.L.A., S.S., T.G., and N.P. performed the experiments. G.S., G.A., T.G. and N.P. analysed the data. G.S. and V.S. wrote the manuscript.

## Competing interests

Authors declare that they have no competing interests.

## Materials & Correspondence

Correspondence and requests for materials should be addressed to Victor Sourjik.

## Extended Data

**Extended Data Fig. 1.**
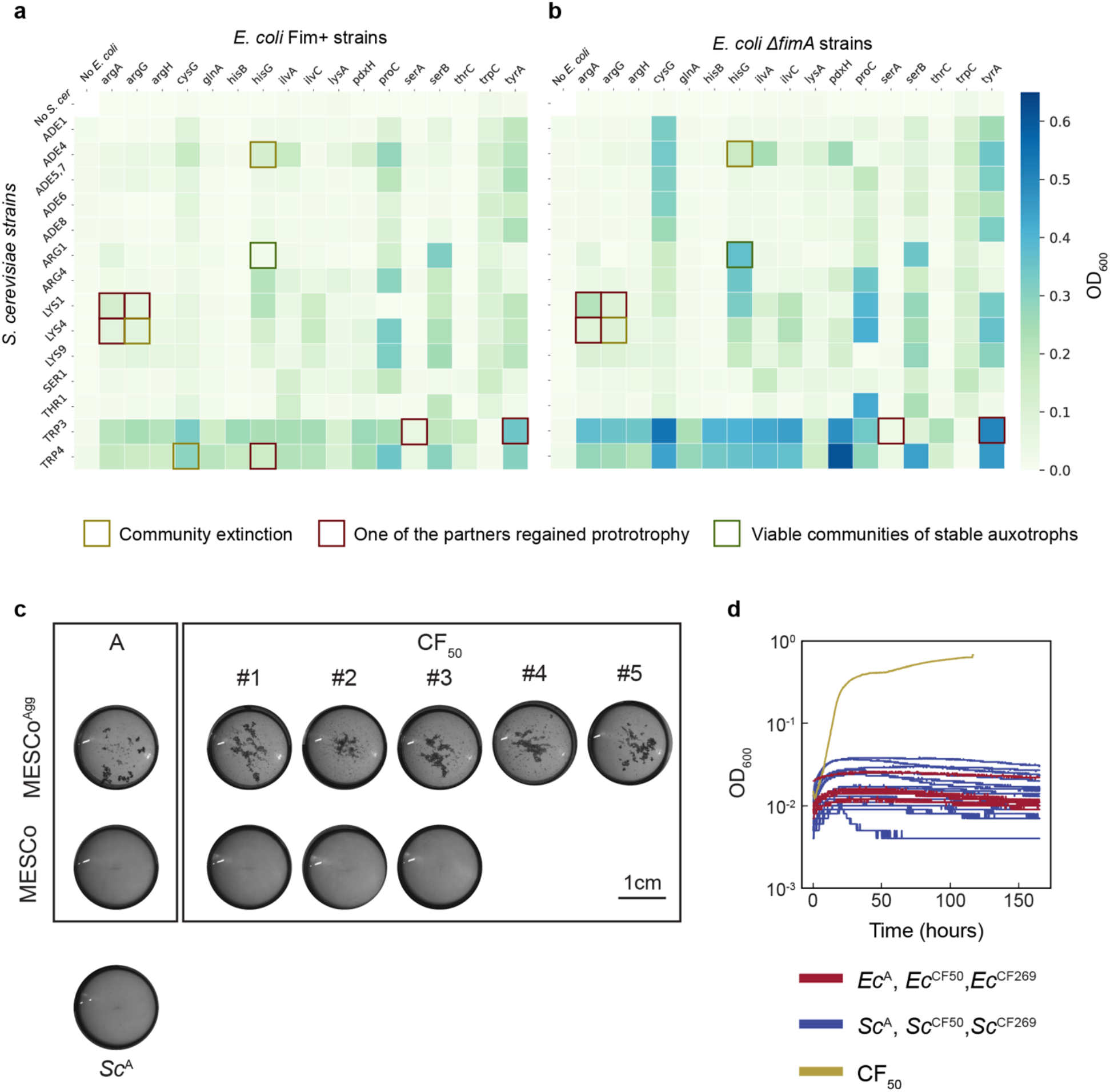
Design and evolution of MESCo communities. (**a,b**) Growth of pairwise co-cultures between different *E. coli* and *S. cerevisiae* strains carrying indicated deletions of metabolic genes, assembled using either fimbriated (Fim^+^; a) or a fimbrialess (fimA; b) *E. coli* partner strains. Color scale indicates density of the co-culture (OD600) after 120 h of cultivation at 30 °C and 200 r.p.m. in CF-MM. Squares indicate communities that have been co-cultured over multiple passages (between 10 and 15), with different colors illustrating their stability and auxotrophy maintenance, as indicated. (**c**) Aggregation test for the non-aggregative (MESCo) and the aggregating (MESCo^Agg^) communities, either ancestral (A) or after co-culture evolution for 50 generations in CF-MM (CF50). To test their aggregation, the communities were incubated in PBS for with shaking at 200 r.p.m. for 1 h at 30 °C in a 24-well plate (Greiner Bio-One GmbH). Also shown is the control culture of the ancestral yeast partner (Sc^A^) alone. (**d**) Growth, measured as OD600 using a plate reader, of monocultures of indicated strains in CF-MM.

**Extended Data Fig. 2.**
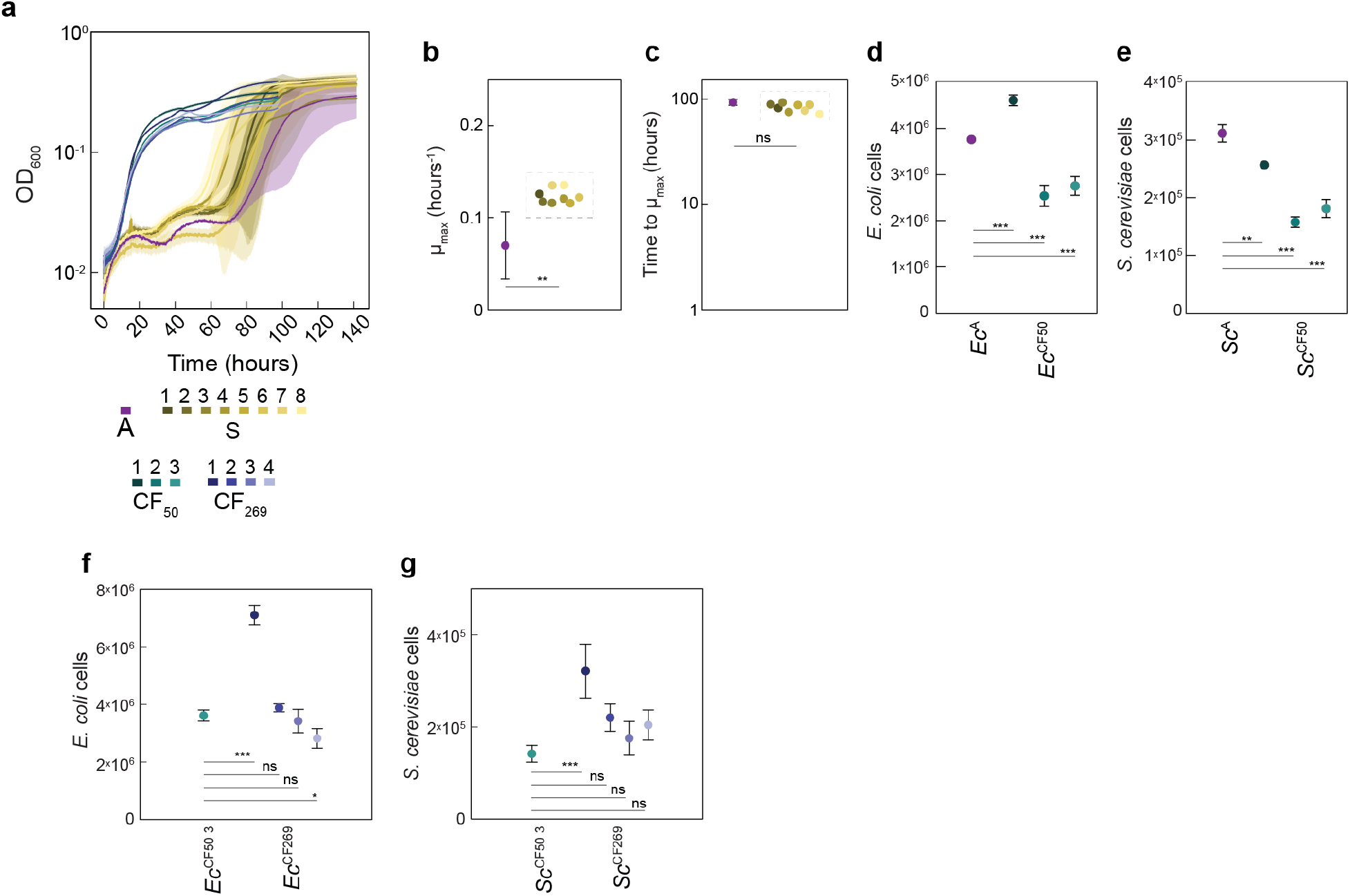
Characterization of growth parameters of the evolved communities. **(a)** Growth profiles of all the eight MESCo communities evolved in AH-MM compared to communities shown in **Figure 1b**. (**b,c**) Maximum growth rate (b) and time to reach it (c) for the ancestral MESCo (A) and the MESCo evolved in AH-MM (S), calculated from the curves shown in (a). The value for A represents the mean from three independent cultures, with the error bar indicating SD. Different colors used for S represent different individual evolved cultures. (**d,e**) Cell counts of *E. coli* (d) and *S. cerevisiae* (e) measured using flow cytometry at the final time point (144 h) of the curves shown in **Figure 1c**. The different colors used for the evolved organisms indicate individual evolved lines. Mean values of n = 3 biological replicates ± SD are shown. (**f,g**) Cell counts of *E. coli* (f) and *S. cerevisiae* (g) for the indicated MESCo communities, measured at the final time point of the curves shown **Figure 1b**. Different colors for the evolved organisms indicate individual evolved lines. Mean of n = 3 biological replicates ± SD. *p* values (ns = *p* > 0.05, * = *p* < 0.05, ** = *p* < 0.01, *** = *p* < 0.001) reported in (b) and (c) are from a two tailed *t*-test assuming unequal variance between the samples while in (d,e,f,g) from a one-way ANOVA followed by Tukey *post-hoc* test.

**Extended Data Fig. 3.**
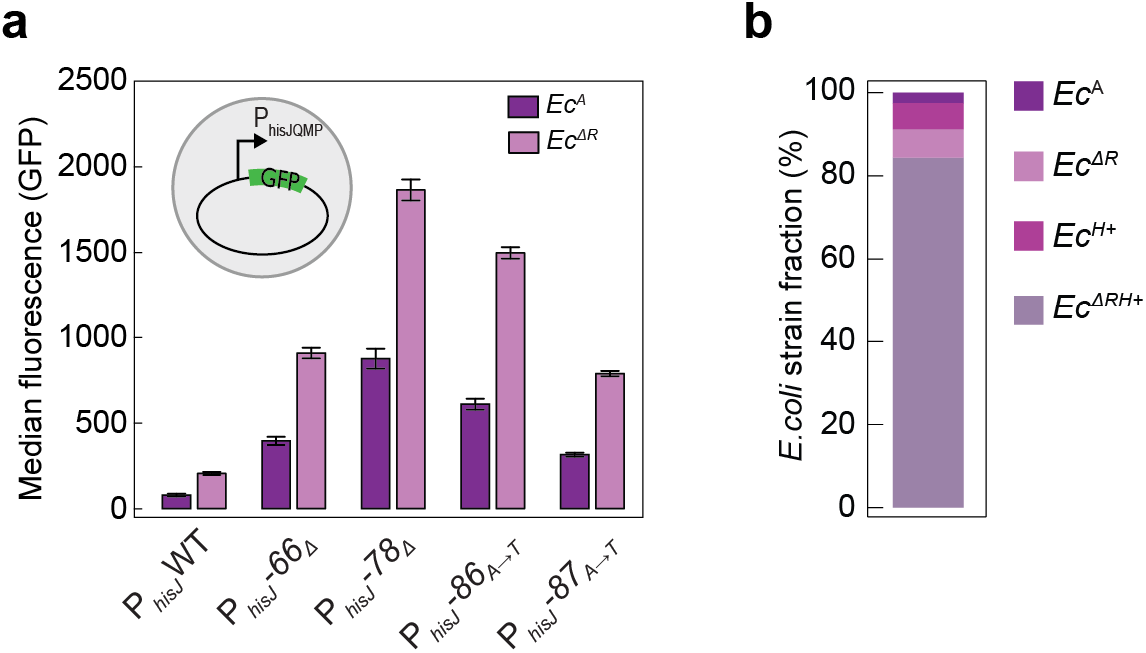
Impact of mutations on *E. coli* and on its fitness. **(a)** Median fluorescence intensity of GFP reporter plasmid, measured using flow cytometry for reporters carrying point mutations identified in the promoter of the *hisJQMP* operon in evolved *E. coli* lines. Reporter plasmids were transformed into ancestral (*Ec*^A^) or *ΔargR* mutant (*Ec^ΔR^*) *E.coli* strain, as indicated, and cultures were grown in CF-MM supplemented with 20 mg/l histidine for 45 h. Mean values of *n* = 3 biological replicates ± SD are shown. (**b**) Average final fraction of indicated *E. coli* strains, initially co-inoculated at the same initial density (OD600 = 0.0125 each) together with the ancestral *S. cerevisiae* strain, and grown in CF-MM for 72h. Mean values of n = 6 biological replicates are shown, with SD (not shown) below 1%.

**Extended Data Fig. 4.**
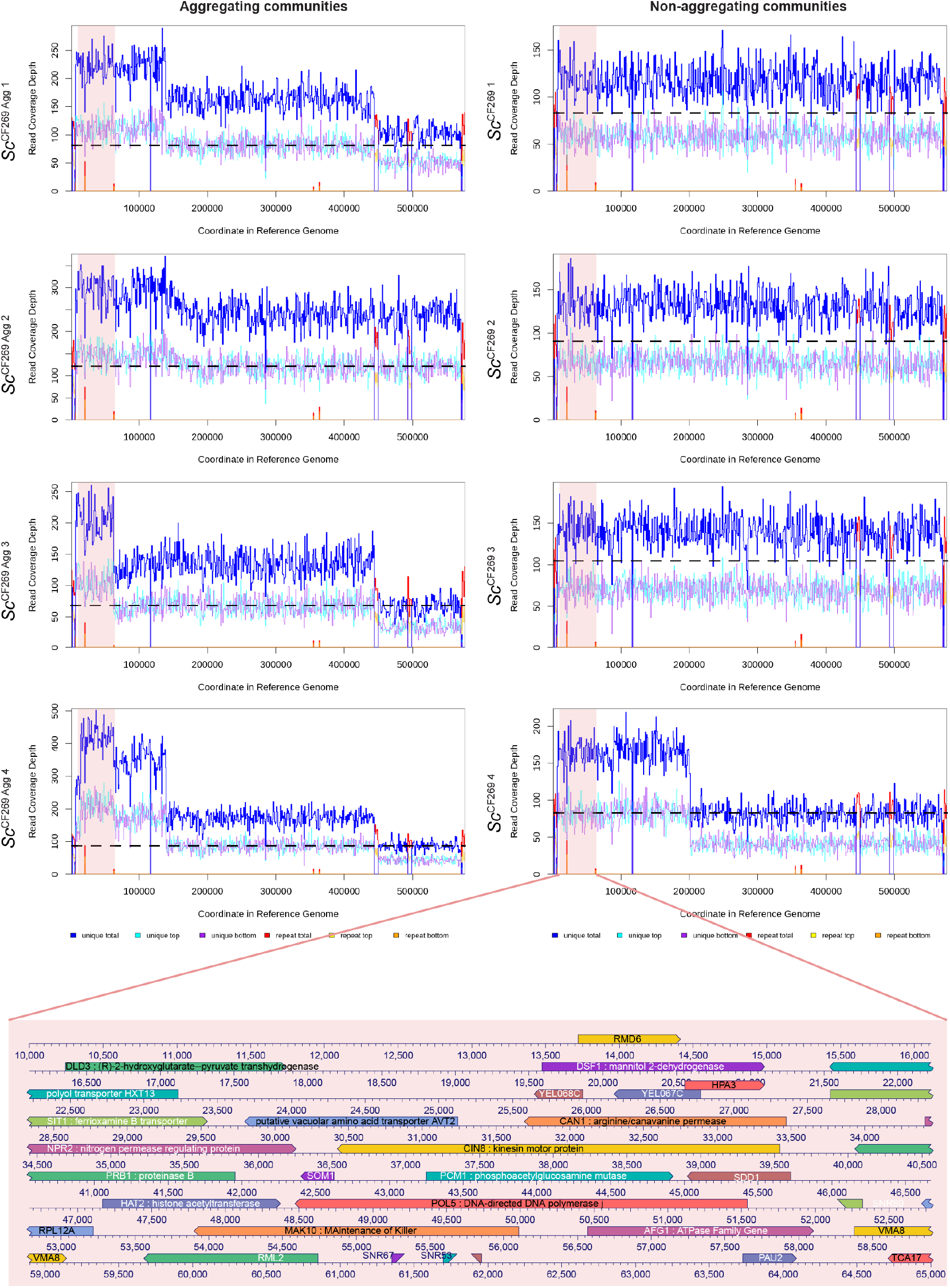
Duplications and aneuploidies in chromosome V in the evolved yeast lines. Read coverage depth obtained in Illumina sequencing on chromosome V in yeast lines in aggregating (left) or non-aggregating (right) MESCo communities evolved for 269 generations in CF-MM. Light blue and purple lines represent the read coverage from paired-end reads matching only once in the reference genome, with the dark blue line indicating their sum. Yellow and orange lines represent the read coverage from pair ends reads matching more than once in the reference genome (e.g. repetitions) and normalized by the number of repetitive sequences found in the genome, with the red line representing their sum. The pink area represents the chromosomal region between nucleotides 10 000 and 60 000 that underwent repetitive events of duplications during evolution, with genes present in this region shown in the area. The dashed black line indicates the mean total read coverage depth for the other chromosomes in each yeast line.

**Extended Data Fig. 5.**
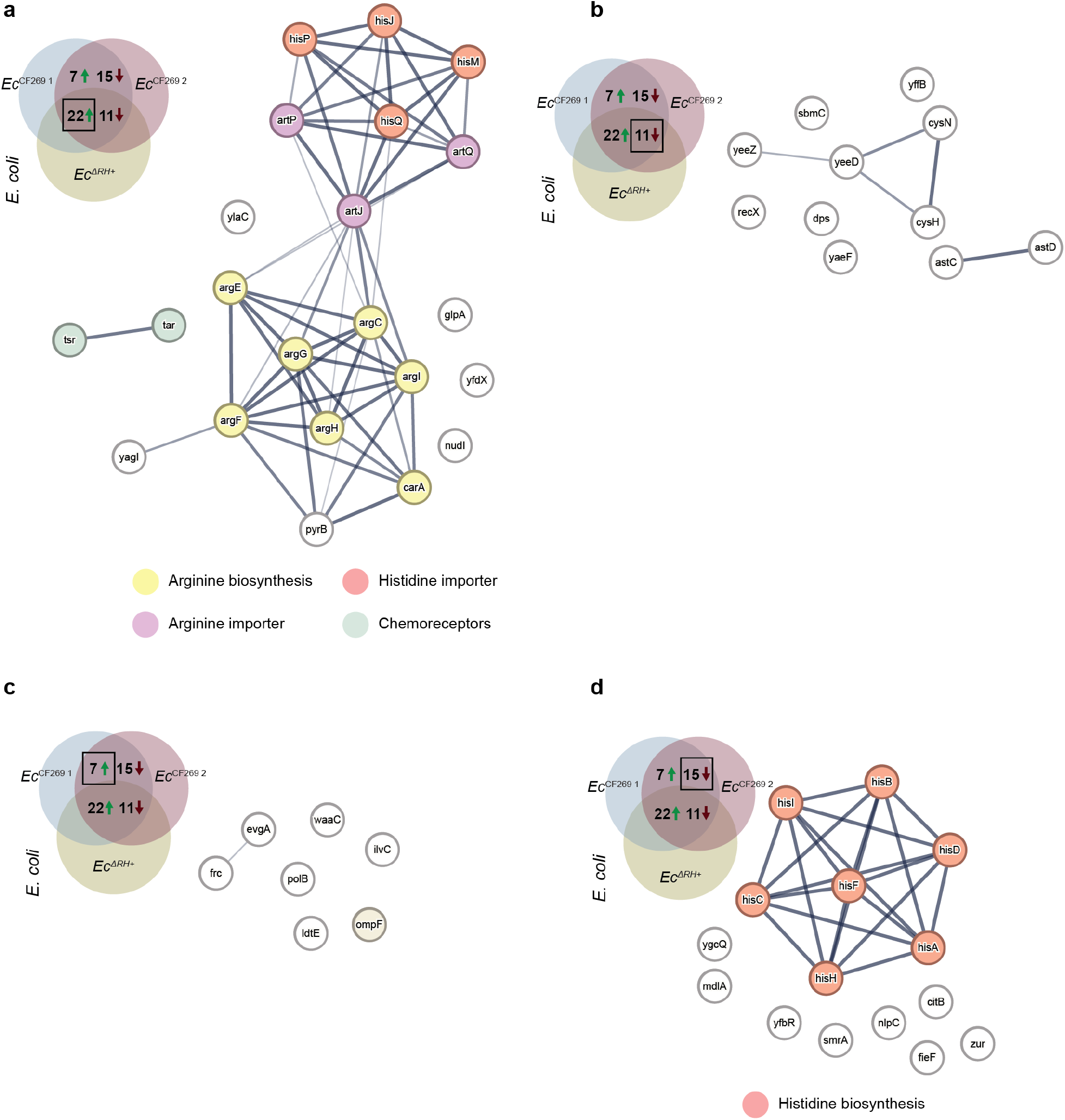
Proteins with different expression levels between indicated *E. coli* strains and the ancestral strain. (**a-d**) STRING^64^ analysis depicting proteins that are upregulated (a,c) (log2(FC) > 1, −Log(p) > 1.5) or downregulated (b,d) (log2(FC) < 1, −Log(p) > 1.5) either in both the mutant and two of the evolved *E. coli* (a,b) or only in the evolved *E. coli* lines (c,d) compared to the ancestral strain. Highlighted are clusters of proteins sharing common functions. Comparison was performed between the evolved communities, the reconstituted communities of mutants carrying major mutations, and the ancestral community. Because of differences in growth between the ancestral and the evolved or mutant communities, only proteins with different expression levels at both 36 h and 100 h were selected. FC = Fold change in total protein intensities.

**Extended Data Fig. 6.**
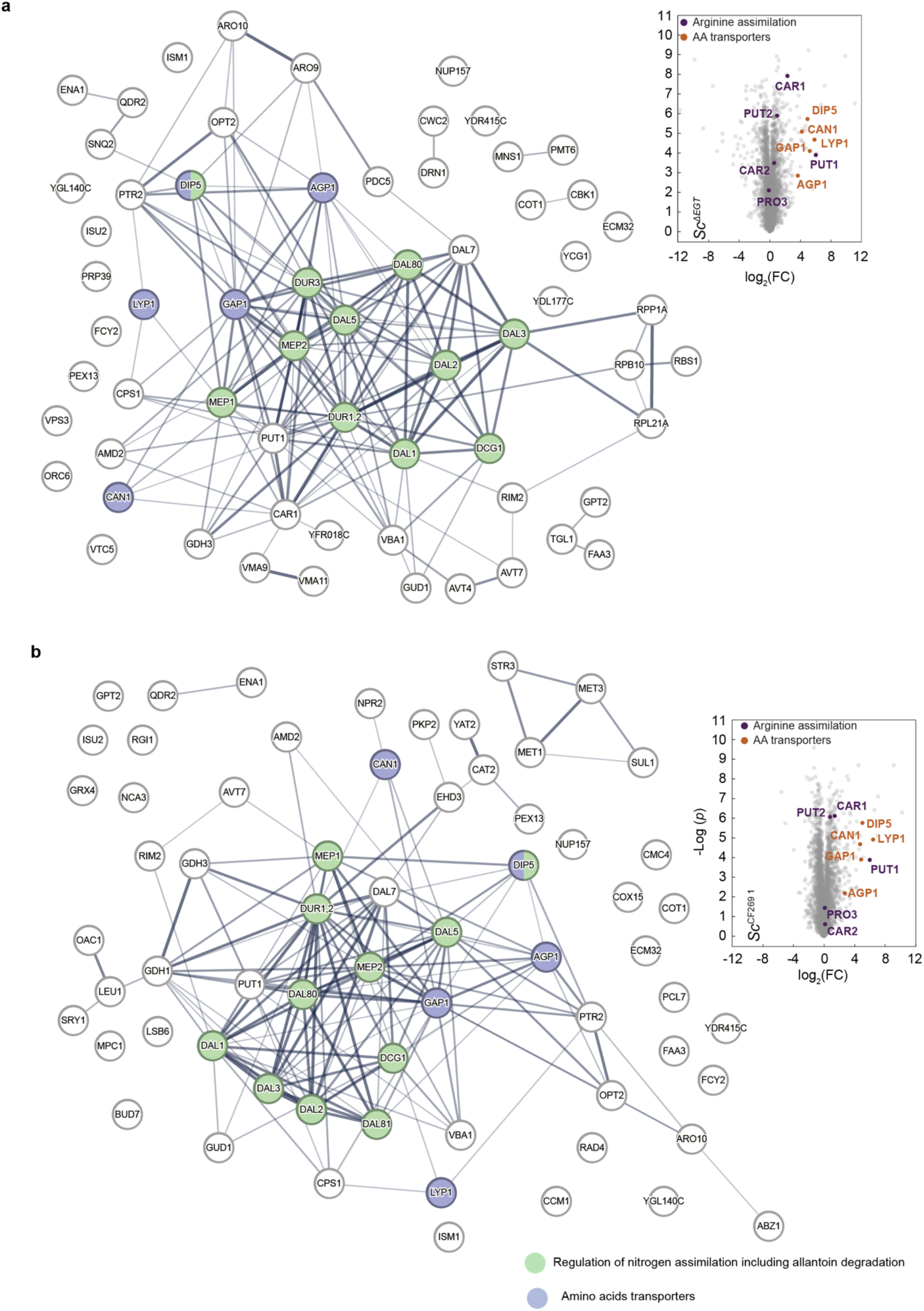
Proteins upregulated in different *S. cereevisiae* lines compared to the ancestral strain in co-cultures with *E. coli ΔargR*. (**a,b**) Volcano plots representing the proteome difference from the ancestral strain and STRING^64^ analysis depicting proteins upregulated (log2(FC) > 2, −Log(*p*) > 2) compared to the ancestral yeast strain in the *Sc^ΔEGT^* mutant (a) and in one of the evolved yeast line (*Sc*^269 1^) (b). All strains were co-cultured with the *Ec^ΔR^* mutant strain for 36 h in CF-MM. For STRING analysis, highlighted are the clusters of proteins involved in either amino acids uptake (blue) or in the regulation of nitrogen utilization and allantoin degradation (green). In volcano plots, the enzymes involved in direct ammonium assimilation (purple) and the amino acid transporters regulated by *ecm21* (orange) are highlighted.

**Extended Data Fig. 7.**
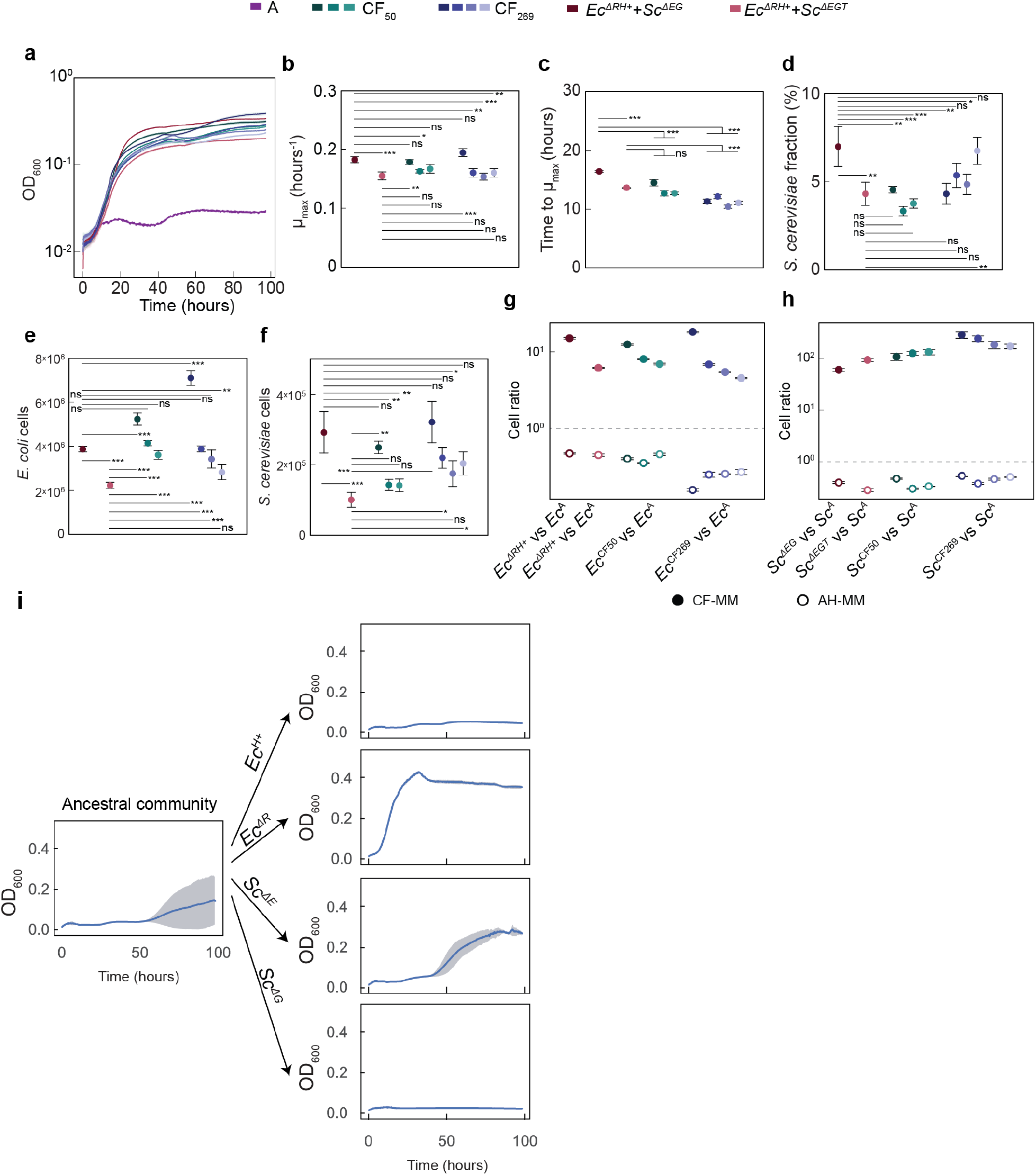
Assessing how well communities of mutants recapitulate the growth parameters of evolved communities. **(a)** Growth of communities of ancestral and mutant *E. coli* and *S. cerevisiae* strains in CF-MM, in comparison with the lines evolved for 50 and 269 generations, as shown in **Figure 1b**. Mean values of *n* = 3 biological replicates *±* SD are shown. (**b-f**) Maximum growth rate (b), time to reach it (c), yeast cell fraction (d), the *E. coli* cell count (e) and the yeast cell count (f) from communities shown in (a) and in **Figure 1b**. Mean values of *n* = 3 biological replicates ± SD are shown. (**g,h**) Cell ratios to the ancestral strain calculated as in **Figure 1g,h**, either in CF-MM or in AH-MM, respectively, for the bacterium (g) or the yeast (h) mutants, in comparison with the evolved strains shown in **Figure 1g,h**. Mean values of *n* = 5 biological replicates ± SD are shown. (**i**) Growth of the community assembled with the ancestral strains, compared to the communities in which one of the organisms is replaced by a strain carrying one of the high-frequency mutations. Mean values of n = 3 biological replicates ± SD are shown. *p* values (ns = *p* > 0.05, * = *p* < 0.05, ** = *p* < 0.01, *** = *p* < 0.001) are from a one-way ANOVA followed by Tukey post-hoc test.

**Extended Data Fig. 8.**
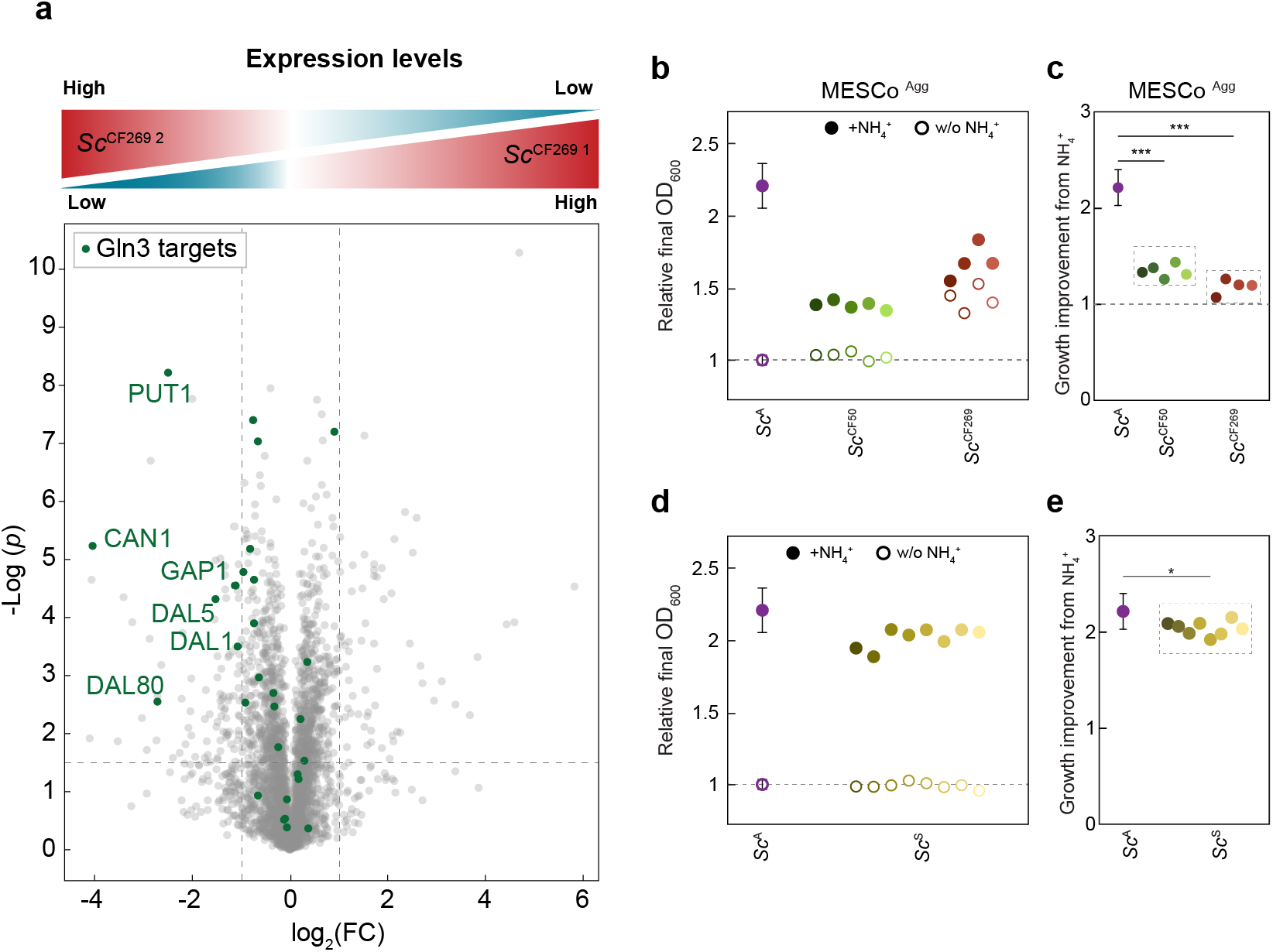
Impact of Gln3 truncation on expression of its targets and effect of ammonium on growth of communities evolved under aggregation or in presence of supplements. **(a)** Volcano plot comparing proteome of the two evolved yeast lines, *Sc*^CF269 1^ and *Sc*^CF269 2^ grown in CF-MM for 36 h. *Sc*^CF269 2^ carries mutation leading to expression of the truncated version of Gln3, whereas *Sc*^CF269 1^ has the full-length variant of Gln3. Mean values of *n* = 4 biological replicates are shown. Known targets of Gln3 are highlighted in green. (**b,c**) Final OD600 of the yeast lines originating from evolved MESCo^Agg^ communities, grown in CF-MM with arginine and either with or without ammonium, relative to the final OD600 of ancestral yeast strain grown in absence of ammonium. (d,e) Relative final OD600 for yeast strains isolated from MESCo communities evolved for 100 generations in AH-MM and grown in CF-MM with arginine either with or without ammonium. In each panel, values represent the average from two biological replicates, except for the Sc^A^ (same as **Figure 3d**) where 11 biological replicates where averaged. Error bars for Sc^A^ represent SD. *p* values (ns = *p* > 0.05, * = *p* < 0.05, ** = *p* < 0.01, *** = *p* < 0.001) are in (c) from a one-way ANOVA followed by Tukey *post-hoc* test while in (e) are from a two tailed *t*-test assuming unequal variance between the samples.

**Extended Data Fig. 9.**
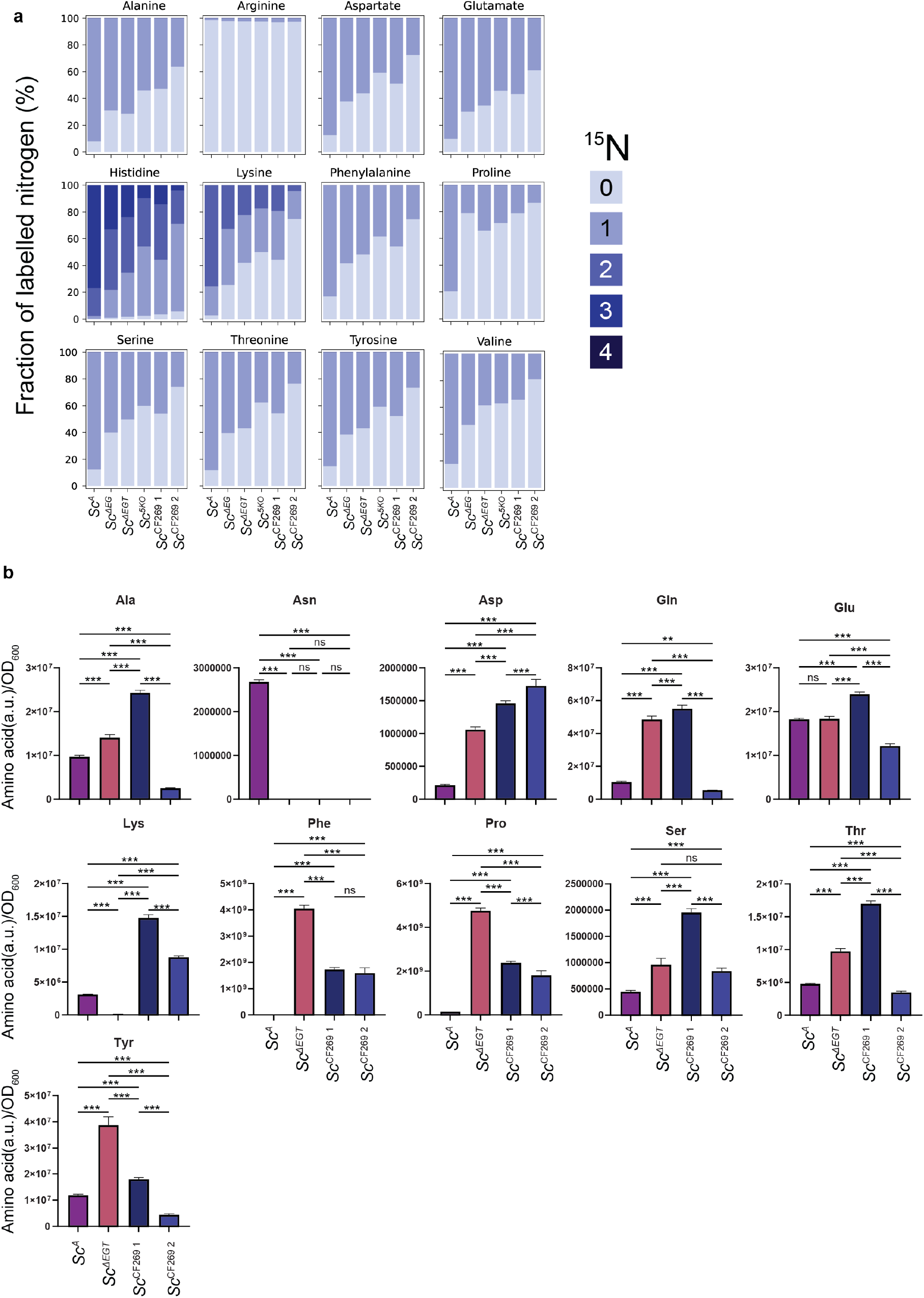
Isotope labeling patters and amino acids exometabolome profiles in indicated yeast strains. **(a)** Average fraction of ^15^N-labelled atoms detected for the full set of proteinogenic amino acids measured in samples described in **Figure 4e,f**. Mean values of *n* = 4 biological replicates are shown, with SD values (not shown) below 3% for all samples except alanine (4%) and valine (13%) for the *Sc^ΔEG^* strain. (**b**) Levels of indicated amino acids expressed in arbitrary units (a.u.), measured as area under the peak from LC-MS measurements of the culture supernatant and normalized to OD600. Cultures were the same as in **Figure 4f** and (a). Mean values of *n* = 4 biological replicates ± SD are shown. *p* values (ns = *p* > 0.05, * = *p* < 0.05, ** = *p* < 0.01, *** = *p* < 0.001) are from a one-way ANOVA followed by Tukey post-hoc test.

**Extended Data Fig. 10.**
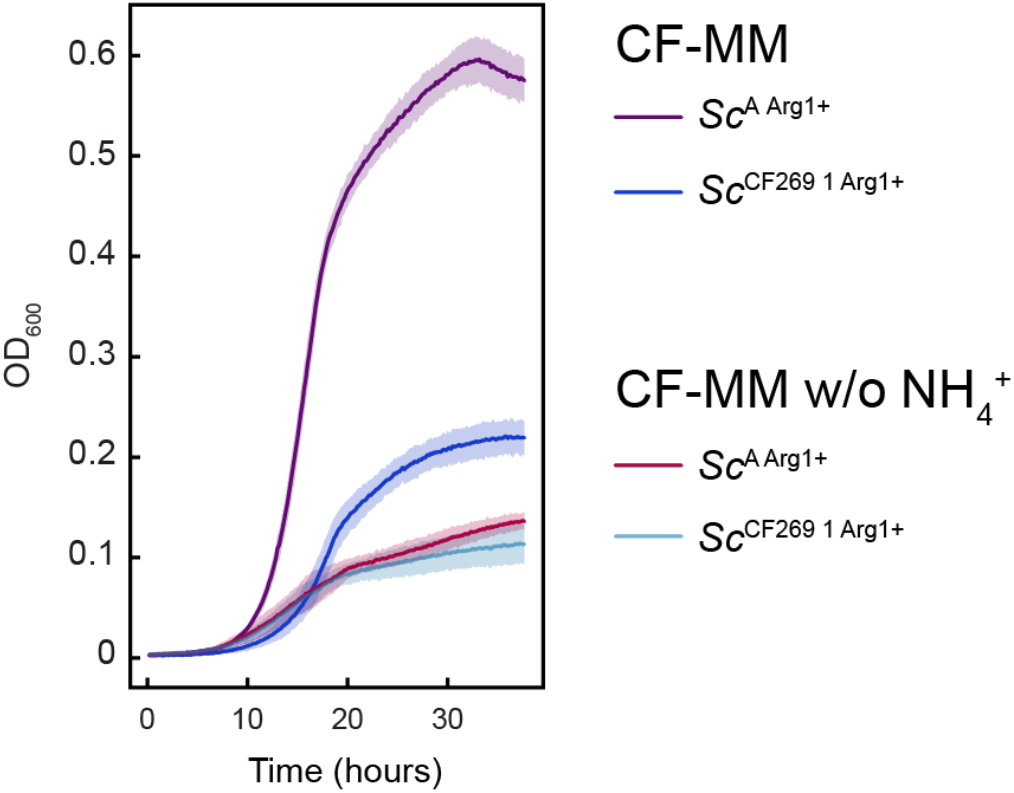
Growth of *S. cerevisiae* in CF-MM with or without ammonium. The ancestral and one of the evolved yeast strains with restored arginine prototrophy grown in CF-MM with or without ammonium. Mean values of *n* = 3 biological replicates ± SD are shown.

## Notes

### Competing Interest Statement

The authors have declared no competing interest.

