## Supplementary Information for "Enhanced metabolic entanglement emerges during the evolution of an interkingdom microbial community"

### **Supplementary Table 1.** Recurrent mutations in *E. coli* evolved lines

**Supplementary Table 1**. Recurrent mutations in *Escherichia coli*

From communities evolved in CF-MM for 50 generations ($CF_{50}$)

| Predicted mutations | | MESCo $^{CF50\ Agg}$ | | | | | MESCo $^{CF50}$ | | | | | |
|---|---|---|---|---|---|---|---|---|---|---|---|---|
| position | mutation | *Ec* $^{50\ 1\ Agg}$ | *Ec* $^{50\ 2\ Agg}$ | *Ec* $^{50\ 3\ Agg}$ | *Ec* $^{50\ 4\ Agg}$ | *Ec* $^{50\ 5\ Agg}$ | *Ec* $^{50\ 1}$ | *Ec* $^{50\ 2}$ | *Ec* $^{50\ 3}$ | annotation | gene | description |
| 982.560 | T→A | | | | | | | 100% | | intergenic (-122/+481) | ompF ← / ← asnS | outer membrane porin /asparaginyl tRNA synthetase |
| 982.560 | T→C | | | | | | 37,50% | | | intergenic (-122/+481) | | |
| 2.420.333 | Δ1 bp | | 100% | | | | | | | intergenic (-66/+155) | hisJ ← / ← argT | histidine-lysine-arginine-ornithine transporter subunit(hisJ) /lysine-arginine-ornithine transporter subunit (argT) |
| 2.420.345 | Δ1 bp | 100% | | | | | | | | intergenic (-78/+143) | | |
| 2.420.353 | A→T | | | | | | 100% | | | intergenic (-86/+135) | | |
| 2.420.354 | A→T | | | | | | | 41,80% | | intergenic (-87/+134) | | |
| 3.378.075 | IS5 (+) +4 bp | | | | | 56% | | | | coding (14-17/471 nt) | argR → | l-arginine-responsive arginine metabolism regulon transcriptional regulator |
| 3.378.089 | IS5 (+) +4 bp | | | | | | | 100% | | coding (28-31/471 nt) | | |
| 3.378.093 | IS5 (+) Δ17 bp | | | | | | | | 63% | coding (32-48/471 nt) | | |
| 3.378.095 | IS | | | | 81% | 93% | | | | coding (34-42/471 nt) | | |
| 3.378.114 | IS5 (–) +4 bp | | | 22% | | | | | | coding (53-56/471 nt) | | |
| 3.378.124 | IS | | | | | | 100% | | | coding (63/471 nt) | | |
| 3.378.137 | C→T | 100% | | | | | | | | Q26* (CAG→TAG) | | |
| 3.378.162 | A→C | | | 17% | | | | | | Q34P (CAG→CCG) | | |
| 3.378.188 | C→T | | | 64% | | | | | | Q43* (CAG→TAG) | | |
| 3.378.387 | C→T | | 100% | | | | | | | A109V (GCT→GTT) | | |
| 3.378.506 | IS5 (–) +4 bp | | | | | | | | 17% | coding (445-448/471 nt) | | |

Communities in red were further evolved in CF-MM up to 269 generations

From communities evolved in CF-MM for 269 generations ($CF_{269}$)

| Predicted mutations | | MESCo$^{269\ Agg}$ (Evolved from Ec$^{50\ 4\ Agg}$) | | | | MESCo$^{269}$ (Evolved from Ec$^{50\ 3}$) | | | | | | |
|---|---|---|---|---|---|---|---|---|---|---|---|---|
| position | mutation | *Ec* $^{269\ 1\ Agg}$ | *Ec* $^{269\ 2\ Agg}$ | *Ec* $^{269\ 3\ Agg}$ | *Ec* $^{269\ 4\ Agg}$ | *Ec* $^{269\ 1}$ | *Ec* $^{269\ 2}$ | *Ec* $^{269\ 3}$ | *Ec* $^{269\ 4}$ | annotation | gene | description |
| 982.560 | T→A | 46% | | | | | 100% | 100% | 100% | intergenic (-122/+481) | ompF ← / ← asnS | outer membrane porin /asparaginyl tRNA synthetase |
| 982.560 | T→G | | 86% | 100% | 100% | | | | | intergenic (-122/+481) | | |
| 982.566 | C→A | | | | | 67% | | | | intergenic (-128/+475) | | |
| 2.420.328 | G→A | 100% | 100% | 100% | 100% | | | | | intergenic (-61/+160) | hisJ ← / ← argT | histidine-lysine-arginine-ornithine transporter subunit(hisJ) /lysine-arginine-ornithine transporter subunit (argT) |
| 2.420.338 | Δ1 bp | | | | | 100% | 100% | 100% | 100% | intergenic (-71/+150) | | |
| 2.420.353 | A→C | | | | | 34% | | | | intergenic (-86/+135) | | |
| 3.378.069 | IS | | | | | 100% | 100% | 100% | 100% | coding (8-16/471 nt)[a] | argR → | l-arginine-responsive arginine metabolism regulon transcriptional regulator |
| 3.378.461 | A→C | 100% | 100% | 100% | 100% | | | | | T134P (ACC→CCC) [a] | | |

* = stop codons
IS = Insertion sequence
(-) (+) = strand orientation
[a]= these mutations replaced the ones in the parental $CF_{50}$ population

### **Supplementary Table 2.** Recurrent mutations in *S. cerevisiae* evolved lines

**Supplementary Table 2.** Recurrent mutations in *S. cerevisiae*

From communities evolved in CF-MM for 50 generations ($CF_{50}$)

| Predicted mutations | | | MESCo$^{CF50\ Agg}$ | | | | | MESCo$^{CF50}$ | | | | | |
|---|---|---|---|---|---|---|---|---|---|---|---|---|---|
| Chromosome | position | mutation | *Sc* $^{50\ 1\ Agg}$ | *Sc* $^{50\ 2\ Agg}$ | *Sc* $^{50\ 3\ Agg}$ | *Sc* $^{50\ 4\ Agg}$ | *Sc* $^{50\ 5\ Agg}$ | *Sc* $^{50\ 1}$ | *Sc* $^{50\ 2}$ | *Sc* $^{50\ 3}$ | annotation | gene | description |
| Chr XV | 1041740 | C→G | 57%** | | 67% | | | | | | A435P (GCT→CCT) | | |
| Chr XV | 1041743 | C→G | | | | 56% | | | | | G434R (GGT→CGT) | | |
| Chr XV | 1041801 | A→T | | | | | | | | 83% | C414* (TGT→TGA) | | |
| Chr XV | 1041945 | C→T | | | | | | | 80% | | W366* (TGG→TGA) | | |
| Chr XV | 1042027 | G→A | | | | | | 7% | | | S339F (TCC→TTC) | | |
| Chr XV | 1042030 | C→T | | | | | 100% | | | | G338D (GGT→GAT) | GDH1 ← | glutamate dehydrogenase |
| Chr XV | 1042465 | G→T | | | | | | | | 20% | T193N (ACT→AAT) | | |
| Chr XV | 1042486 | G→A | | | | | | 100% | | | S186F (TCT→TTT) | | |
| Chr XV | 1042811 | G→A | | | | | | | 28% | | R78C (CGT→TGT) | | |
| Chr XV | 1042891 | Δ1 bp | | 100% | | | | | | | coding (152/1365 nt) | | |
| Chr XV | 1042942 | Δ1 bp | | | | 37% | | | | | coding (101/1365 nt) | | |
| Chr XV | 1042945 | A→C | | | | 37% | | | | | L33W (TTG→TGG) | | |
| Chr II | 25.826 | Δ1 bp | 39,20% | 25,60% | 31,50% | | | | | | coding (2474/3354 nt) | | |
| Chr II | 25.865 | Δ1 bp | | | | | | | | 100% | coding (2435/3354 nt) | | |
| Chr II | 25.937 | Δ1 bp | | | | | | 12,70% | | | coding (2363/3354 nt) | | |
| Chr II | 26.262 | C→A | 38,60% | 13,10% | 56,60% | | | | | | E680* (GAG→TAG) | | |
| Chr II | 26.326 | A→C | | | | 46,70% | | | | | Y658* (TAT→TAG) | | |
| Chr II | 26,394:1 | +A | | | | | | | 13,50% | | coding (1906/3354 nt) | | |
| Chr II | 26.401 | T→G | | | | | | | 13,20% | | T633T (ACA→ACC) | | |
| Chr II | 26.465 | A→T | | | | | | | | 18,30% | L612* (TTG→TAG) | ECM21 ← | Alpha-arrestin, ubiquitin ligase adaptor |
| Chr II | 26.529 | A→C | | | | | 89,00% | | | | Y591D (TAC→GAC) | | |
| Chr II | 26.651 | G→C | | | | | | | 72,80% | | S550* (TCA→TGA) | | |
| Chr II | 26.922 | C→A | | | | | | 12,70% | | | E460* (GAG→TAG) | | |
| Chr II | 27.002 | Δ50 bp | | | | | | | | 16,60% | coding (1249-1298/3354 nt) | | |
| Chr II | 27.057 | C→A | | | | 16,90% | | | | | E415* (GAG→TAG) | | |
| Chr II | 27.185 | G→T | | | | 35,30% | | | | | S372* (TCA→TAA) | | |
| Chr II | 27.690 | T→A | | | | | | 64,70% | | | R204* (AGA→TGA) | | |
| Chr IV | 152.361 | A→G | | | 9,90% | | | | | | S1094P (TCT→CCT) | | |
| Chr IV | 153.790 | C→T | | | | | | | | 26,00% | W617* (TGG→TGA) ‡ | GLT1 ← | glutamate synthase |
| Chr IV | 153.791 | C→T | | | | | | | | 25,90% | W617* (TGG→TAG) ‡ | | |
| Chr IV | 153.902 | C→G | | | | | | | 66,80% | 22,20% | R580P (CGT→CCT) | | |
| Chr VII | 733.110 | Δ1 bp | | | | | 100% | | | | intergenic (-183/-302) | MEP1 ← / → YGR121W-A | ammonium permease /hypothetical protein |
| Chr V | 231.443 | C→G | | | | | | | | 7,10% | S550* (TCA→TGA) | GLN3 → | nitrogen-responsive transcriptional regulator |
| Chr I | 28.316 | C→A | | | | | | | | 10,10% | intergenic (-348/-3251) | FLO9 ← / → GDH3 | flocculin FLO9/glutamate dehydrogenase |

Communities in red were further evolved in CF-MM up to 269 generations

From communities evolved in CF-MM for 269 generations ($CF_{269}$)

| Predicted mutations | | | MESCo$^{269\ Agg}$ (Evolved from Sc$^{50\ 4\ Agg}$) | | | | MESCo$^{269}$ (Evolved from Sc$^{50\ 3}$) | | | | | | |
|---|---|---|---|---|---|---|---|---|---|---|---|---|---|
| Chromosome | position | mutation | *Sc* $^{269\ 1\ Agg}$ | *Sc* $^{269\ 2\ Agg}$ | *Sc* $^{269\ 3\ Agg}$ | *Sc* $^{269\ 4\ Agg}$ | *Sc* $^{269\ 1}$ | *Sc* $^{269\ 2}$ | *Sc* $^{269\ 3}$ | *Sc* $^{269\ 4}$ | annotation | gene | description |
| Chr XV | 1.041.743 | C→G | 27,20% | 100% | 100% | 100% | | | | | G434R (GGT→CGT) | | |
| Chr XV | 1.041.801 | A→T | | | | | 9,00% | 100% | 78,90% | 100% | C414* (TGT→TGA) | | |
| Chr XV | 1.042.465 | G→T | | | | | 90,20% | | 28,10% | | T193N (ACT→AAT) | GDH1 ← | glutamate dehydrogenase |
| Chr XV | 1.042.882 | C→T | | | | | | 6,90% | | | G54D (GGT→GAT) | | |
| Chr XV | 1.042.943 | Δ1 bp | 72,40% | | | | | | | | coding (100/1365 nt) | | |
| Chr XV | 1.042.945 | A→C | 76,50% | | | | | | | | L33W (TTG→TGG) | | |
| Chr II | 25.867 | Δ1 bp | | | | | | | 100% | 100% | coding (2433/3354 nt) | | |
| Chr II | 26.326 | A→C | 24,30% | 100% | 100% | 100% | | | | | Y658* (TAT→TAG) | | |
| Chr II | 26.465 | A→T | | | | | 74,80% | | 24,30% | | L612* (TTG→TAG) [a] | ECM21 ← | Alpha-arrestin, ubiquitin ligase adaptor |
| Chr II | 27.002 | Δ50 bp | | | | | | 77,90% | | | coding (1249-1298/3354 nt) [a] | | |
| Chr II | 27.185 | G→T | 66,10% | | | | | | | | S372* (TCA→TAA) | | |
| Chr IV | 149.315 | C→T | | 11,00% | | | | | | | C2109Y (TGT→TAT) | | |
| Chr IV | 149.837 | T→C | | | | 7,40% | | | | | D1935G (GAT→GGT) | | |
| Chr IV | 150.116 | A→T | | | | 28,80% | | | | | L1842* (TTG→TAG) | | |
| Chr IV | 150.422 | C→G | | | | 9,50% | | | | | C1740S (TGT→TCT) | | |
| Chr IV | 150.803 | A→T | | 65,40% | | | | | | | L1613* (TTG→TAG) | | |
| Chr IV | 150.972 | C→A | 20,80% | | | | | | | | E1557* (GAG→TAG) | GLT1 ← | glutamate synthase |
| Chr IV | 153.790 | 2 bp→TT | | | | | | | | 100% | coding (1850-1851/6438 nt) [a] | | |
| Chr IV | 153.790 | C→T | | | | | 18,50% | 24,70% | 70,40% | | W617* (TGG→TGA) ‡ | | |
| Chr IV | 153.791 | C→T | | | | | 18,40% | 24,70% | 70,40% | | W617* (TGG→TAG) ‡ | | |
| Chr IV | 153.902 | C→G | | | | | | 73,50% | | | R580P (CGT→CCT) | | |
| Chr IV | 154.617 | C→A | | | | 6,80% | | | | | E342* (GAA→TAA) | | |
| Chr VII | 731.823 | G→C | | 62,40% | | | | | | | Q369E (CAA→GAA) | | |
| Chr VII | 732.008 | A→G | | | | | | | 25,90% | | V307A (GTG→GCG) | | |
| Chr VII | 732.027 | C→T | | | | | | 70,40% | | | G301S (GGC→AGC) | | |
| Chr VII | 732.038 | G→A | | | | | | | | 100% | S297L (TCA→TTA) | MEP1 ← | ammonium permease |
| Chr VII | 732.099 | C→G | | | | | | | 9,40% | | G277R (GGT→CGT) | | |
| Chr VII | 732.779 | A→T | | | | | | | 34,50% | | L50Q (CTA→CAA) | | |
| Chr VII | 732.791 | C→T | 13,80% | | | | | | | | R46K (AGG→AAG) | | |
| BK006949 | 811.036 | C→T | 7,30% | | | | | | | | G473D (GGC→GAC) | | |
| BK006949 | 811.413 | C→T | | | | | | 6,30% | | | W347* (TGG→TGA) | | |
| BK006949 | 811.595 | T→A | | | | | 17,70% | 20,00% | 57,00% | 100% | S287C (AGC→TGC) | MEP3 ← | ammonium permease |
| BK006949 | 811.945 | A→T | | 15,50% | | | | | | | V170E (GTA→GAA) | | |
| BK006949 | 812.185 | G→T | | | | | | 77% | | | S90* (TCA→TAA) | | |
| BK006949 | 812.335 | G→A | | | | | 75% | | | | S40F (TCC→TTC) | | |
| BK006947 | 358.371 | G→A | | | 6% | | | | | | A307T (GCC→ACC) | MEP2 → | ammonium permease |
| BK006939 | 231405 | A→G | | 35% | | 100% | | | | | A537A (GCA→GCG) | | |
| BK006939 | 231409 | Δ76 bp | | | | 100% | | | | | coding (1615-1690/2193 nt) | GLN3 → | nitrogen-responsive transcriptional regulator |
| BK006939 | 231443 | C→G | | | | | | | 37% | | S550* (TCA→TGA) | | |
| BK006939 | 231473 | C→A | | | | | | 57% | | | S560* (TCA→TAA) | | |
| BK006935 | 31123 | Δ1 bp | 13% | 8% | | | 16% | 26% | 14% | 11% | intergenic (-3155/-444) | | |
| BK006935 | 31124 | Δ1 bp | 13% | 8% | | | 16% | 26% | 14% | 11% | intergenic (-3156/-443) | | |
| BK006935 | 31126 | +ATAT | | | 18% | | | | | 11% | intergenic (-3158/-441) | | |
| BK006935 | 31122:1 | +A | | 17% | | 13% | | 13% | | | intergenic (-3154/-445) | | |
| BK006935 | 31122:2 | +T | | 17% | | 13% | | 13% | | | intergenic (-3154/-445) | | |
| BK006935 | 31484 | Δ1 bp | | | | | | 10% | | 7% | intergenic (-3516/-83) | FLO9 ← / → GDH3 | flocculin FLO9/glutamate dehydrogenase |
| BK006935 | 31485 | Δ1 bp | | | | | | | | 6% | intergenic (-3517/-82) | | |
| BK006935 | 31504:1 | +A | | | | 13% | | | | 9% | intergenic (-3536/-63) | | |
| BK006935 | 31504:2 | +T | | | | 13% | | | | 9% | intergenic (-3536/-63) | | |
| BK006935 | 31508 | +ATAT | | | | | | | 7% | | intergenic (-3540/-59) | | |
| BK006935 | 31901 | G→A | | 25% | | | | | | | G112D (GGT→GAT) | | |
| BK006935 | 32012 | C→A | | | | | | | 41% | | A149E (GCA→GAA) | | |
| BK006935 | 32164 | T→A | | 16% | | | | | | | Y200N (TAC→AAC) | GDH3 → | glutamate dehydrogenase |
| BK006935 | 32596 | T→C | | | | | | | 8% | | S344P (TCT→CCT) | | |
| BK006935 | 32888 | C→T | | | | | | | | 12% | A441V (GCC→GTC) | | |

* = stop codons

‡ = the mutations are co-occuring in the same strain

** Coverage too low for mutations call via breseq. Subsequent analysis via IGV revealed 24 counts out of a total of 42 as C->G

[a] = these mutations replaced the ones in the parental $CF_{50}$ population

**Supplementary Table 3.** Statistical analysis **Figure 2i**. Compact Letter Display (CLD) from the Tukey test performed as post-hoc analysis. Strains with the same letter have no statistical difference in intensity levels for the indicated protein. The *p* values from the ANOVA analysis are also reported.

| | **Strains** | | | | |
|---|---|---|---|---|---|
| **Protein** | $\textbf{\textit{Sc}}^{A}$ | $\textbf{\textit{Sc}}^{\Delta EG}$ | $\textbf{\textit{Sc}}^{\Delta EGT}$ | $\textbf{\textit{Sc}}^{CF269\ 1}$ | **ANOVA p value** |
| **PUT2** | a | b | c | d | $p<0.001$ |
| **PUT1** | a | b | c | c | $p<0.001$ |
| **CAR2** | a | b | b | a | $p<0.001$ |
| **CAR1** | a | b | c | d | $p<0.001$ |
| **PRO3** | a | a | b | c | $p<0.001$ |
| **CAN1** | a | b | c | c | $p<0.001$ |
| **GAP1** | a | b | c | c | $p<0.001$ |
| **AGP1** | a | a | b | b | $p<0.001$ |
| **DIP5** | a | b | c | c | $p<0.001$ |
| **LYP1** | a | b,c | cd | d | $p<0.001$ |

**Supplementary Table 4.** *Escherichia coli* strains used in this study

| ***Escherichia coli* strains** | **Name** | **Reference** |
|---|---|---|
| *E. coli* BW25113 *ΔhisG:kan*$^R$ | $Ec^{\text{A Agg}}$ | [51,53] |
| *E. coli* BW25113 *ΔfimA:kan*$^R$ | | [51,53] |
| *E. coli* BW25113 *ΔfimA:frt ΔhisG:kan*$^R$ | $Ec^{\text{A}}$ | This work |
| *E. coli* BW25113 *ΔfimA:frt ΔhisG:frt* | | This work |
| *E. coli* BW25113 *ΔfimA:frt ΔhisG:frt ΔargR:frt* | $Ec^{\Delta R}$ | This work |
| *E. coli* BW25113 *ΔfimA:frt ΔhisG:frt* $p_{hisJ}$*-78Δ* | $Ec^{\Delta H+}$ | This work |
| *E. coli* BW25113 *ΔfimA:frt ΔhisG:frt ΔargR:frt* $p_{hisJ}$*-78Δ* | $Ec^{\Delta RH+}$ | This work |
| *E. coli* BW25113 *ΔhisG:kan*$^R$ | $Ec^{\text{CF50 1 Agg}}$ | This work |
| *E. coli* BW25113 *ΔhisG:kan*$^R$ | $Ec^{\text{CF50 2 Agg}}$ | This work |
| *E. coli* BW25113 *ΔhisG:kan*$^R$ | $Ec^{\text{CF50 3 Agg}}$ | This work |
| *E. coli* BW25113 *ΔhisG:kan*$^R$ | $Ec^{\text{CF50 4 Agg}}$ | This work |
| *E. coli* BW25113 *ΔhisG:kan*$^R$ | $Ec^{\text{CF50 5 Agg}}$ | This work |
| *E. coli* BW25113 *ΔhisG:kan*$^R$ | $Ec^{\text{CF269 1 Agg}}$ | This work |
| *E. coli* BW25113 *ΔhisG:kan*$^R$ | $Ec^{\text{CF269 2 Agg}}$ | This work |
| *E. coli* BW25113 *ΔhisG:kan*$^R$ | $Ec^{\text{CF269 3 Agg}}$ | This work |
| *E. coli* BW25113 *ΔhisG:kan*$^R$ | $Ec^{\text{CF269 4 Agg}}$ | This work |
| *E. coli* BW25113 *ΔfimA:frt ΔhisG:frt* | $Ec^{\text{CF50 1}}$ | This work |
| *E. coli* BW25113 *ΔfimA:frt ΔhisG:frt* | $Ec^{\text{CF50 2}}$ | This work |
| *E. coli* BW25113 *ΔfimA:frt ΔhisG:frt* | $Ec^{\text{CF50 3}}$ | This work |
| *E. coli* BW25113 *ΔfimA:frt ΔhisG:frt* | $Ec^{\text{CF269 1}}$ | This work |
| *E. coli* BW25113 *ΔfimA:frt ΔhisG:frt* | $Ec^{\text{CF269 2}}$ | This work |
| *E. coli* BW25113 *ΔfimA:frt ΔhisG:frt* | $Ec^{\text{CF269 3}}$ | This work |
| *E. coli* BW25113 *ΔfimA:frt ΔhisG:frt* | $Ec^{\text{CF269 4}}$ | This work |
| *E. coli* BW25113 *ΔfimA:frt ΔhisG:frt* | $Ec^{\text{S 1}}$ | This work |
| *E. coli* BW25113 *ΔfimA:frt ΔhisG:frt* | $Ec^{\text{S 2}}$ | This work |
| *E. coli* BW25113 *ΔfimA:frt ΔhisG:frt* | $Ec^{\text{S 3}}$ | This work |
| *E. coli* BW25113 *ΔfimA:frt ΔhisG:frt* | $Ec^{\text{S 4}}$ | This work |
| *E. coli* BW25113 *ΔfimA:frt ΔhisG:frt* | $Ec^{\text{S 5}}$ | This work |
| *E. coli* BW25113 *ΔfimA:frt ΔhisG:frt* | $Ec^{\text{S 6}}$ | This work |
| *E. coli* BW25113 *ΔfimA:frt ΔhisG:frt* | $Ec^{\text{S 7}}$ | This work |

| | | |
|---|---|---|
| *E. coli* BW25113 *ΔfimA:frt ΔhisG:frt* | $Ec^{S}$ [8] | This work |
| *E. coli* BW25113 *ΔfimA:frt ΔhisG:hisG* | $Ec^{CF269\ 1\ HisG+}$ | This work |
| *E. coli* BW25113 *ΔfimA:frt ΔhisG:hisG* | $Ec^{CF269\ 2\ HisG+}$ | This work |
| *E. coli* BW25113 *ΔargR:*$kan^R$ | | [51,53] |
| *E. coli* BW25113 *ΔargA:*$kan^R$ | | [51,53] |
| *E. coli* BW25113 *ΔargG:*$kan^R$ | | [51,53] |
| *E. coli* BW25113 *ΔargH:*$kan^R$ | | [51,53] |
| *E. coli* BW25113 *ΔcysG:*$kan^R$ | | [51,53] |
| *E. coli* BW25113 *ΔglnA:*$kan^R$ | | [51,53] |
| *E. coli* BW25113 *ΔhisB:*$kan^R$ | | [51,53] |
| *E. coli* BW25113 *ΔilvA:*$kan^R$ | | [51,53] |
| *E. coli* BW25113 *ΔilvC:*$kan^R$ | | [51,53] |
| *E. coli* BW25113 *ΔlysA:*$kan^R$ | | [51,53] |
| *E. coli* BW25113 *ΔpdxH:*$kan^R$ | | [51,53] |
| *E. coli* BW25113 *ΔproC:*$kan^R$ | | [51,53] |
| *E. coli* BW25113 *ΔserA:*$kan^R$ | | [51,53] |
| *E. coli* BW25113 *ΔserB:*$kan^R$ | | [51,53] |
| *E. coli* BW25113 *ΔthrC:*$kan^R$ | | [51,53] |
| *E. coli* BW25113 *ΔtrpC:*$kan^R$ | | [51,53] |
| *E. coli* BW25113 *ΔtyrA:*$kan^R$ | | [51,53] |
| *E. coli* BW25113 *ΔfimA:frt ΔargA:*$kan^R$ | | This work |
| *E. coli* BW25113 *ΔfimA:frt ΔargG:*$kan^R$ | | This work |
| *E. coli* BW25113 *ΔfimA:frt ΔargH:*$kan^R$ | | This work |
| *E. coli* BW25113 *ΔfimA:frt ΔcysG:*$kan^R$ | | This work |
| *E. coli* BW25113 *ΔfimA:frt ΔglnA:*$kan^R$ | | This work |
| *E. coli* BW25113 *ΔfimA:frt ΔhisB:*$kan^R$ | | This work |
| *E. coli* BW25113 *ΔfimA:frt ΔilvA:*$kan^R$ | | This work |
| *E. coli* BW25113 *ΔfimA:frt ΔilvC:*$kan^R$ | | This work |
| *E. coli* BW25113 *ΔfimA:frt ΔlysA:*$kan^R$ | | This work |
| *E. coli* BW25113 *ΔfimA:frt ΔpdxH:*$kan^R$ | | This work |
| *E. coli* BW25113 *ΔfimA:frt ΔproC:*$kan^R$ | | This work |

| | | |
|---|---|---|
| *E. coli* BW25113 *ΔfimA:frt ΔserA:kan*$^R$ | | This work |
| *E. coli* BW25113 *ΔfimA:frt ΔserB:kan*$^R$ | | This work |
| *E. coli* BW25113 *ΔfimA:frt ΔthrC:kan*$^R$ | | This work |
| *E. coli* BW25113 *ΔfimA:frt ΔtrpC:kan*$^R$ | | This work |
| *E. coli* BW25113 *ΔfimA:frt ΔtyrA:kan*$^R$ | | This work |

**Supplementary Table 5.** *Saccharomyces cerevisiae* strains used in this study

| ***Saccharomyces cerevisiae*** | **Name** | **Reference** |
|---|---|---|
| *S. cerevisiae* BY4741*Δarg1:kanMX*$^{R}$ | | [54] |
| *S. cerevisiae* BY4741*Δhis3::HIS3-Pglk1-mTurquoise2-Tglk1 Δarg1:kanMX*$^{R}$ | *Sc*$^{A}$ (mTurq2) | This work |
| *S. cerevisiae* BY4741*Δhis3::HIS3-Pglk1-mTurquoise2-Tglk1 Δarg1:kanMX*$^{R}$ *Δgdh1:loxP* | *Sc*$^{ΔG}$ (mTurq2) | This work |
| *S. cerevisiae* BY4741*Δhis3::HIS3-Pglk1-mTurquoise2-Tglk1 Δarg1:kanMX*$^{R}$ *Δecm21:loxP* | *Sc*$^{ΔE}$ (mTurq2) | This work |
| *S. cerevisiae* BY4741*Δhis3::HIS3-Pglk1-mTurquoise2-Tglk1 Δarg1:kanMX*$^{R}$ *Δgdh1:loxP Δecm21:loxP* | *Sc*$^{ΔEG}$ (mTurq2) | This work |
| *S. cerevisiae* BY4741*Δhis3::HIS3-Pglk1-mTurquoise2-Tglk1 Δarg1:kanMX*$^{R}$ *Δgdh1:loxP Δecm21:loxP Δglt1:loxP* | *Sc*$^{ΔEGT}$ (mTurq2) | This work |
| *S. cerevisiae* BY4741*Δhis3::HIS3-Pglk1-mTurquoise2-Tglk1 Δarg1:kanMX*$^{R}$ *Δgdh1:loxP Δecm21:loxP Δglt1:loxP Δgdh3:loxP Δgdh2:HyB* | *Sc*$^{5KO}$ (mTurq2) | This work |
| *S. cerevisiae* BY4741*Δhis3::HIS3-Pglk1-mNeonGreen-Tglk1 Δarg1:kanMX*$^{R}$ | *Sc*$^{A}$ (mNeGr) | This work |
| *S. cerevisiae* BY4741*Δhis3::HIS3-Pglk1-mNeonGreen-Tglk1 Δarg1:kanMX*$^{R}$ *Δgdh1:loxP* | *Sc*$^{ΔG}$ (mNeGr) | This work |
| *S. cerevisiae* BY4741*Δhis3::HIS3-Pglk1-mNeonGreen-Tglk1 Δarg1:kanMX*$^{R}$ *Δgdh1:loxP Δecm21:loxP* | *Sc*$^{ΔEG}$ (mNeGr) | This work |
| *S. cerevisiae* BY4741*Δhis3::HIS3-Pglk1-mNeonGreen-Tglk1 Δarg1:kanMX*$^{R}$ *Δgdh1:loxP Δecm21:loxP Δglt1:loxP* | *Sc*$^{ΔEGT}$ (mNeGr) | This work |
| *S. cerevisiae* BY4741*Δhis3::HIS3-Pglk1-mTurquoise2-Tglk1 Δarg1:kanMX*$^{R}$ | *Sc*$^{CF50\ 1}$ | This work |
| *S. cerevisiae* BY4741*Δhis3::HIS3-Pglk1-mTurquoise2-Tglk1 Δarg1:kanMX*$^{R}$ | *Sc*$^{CF50\ 2}$ | This work |
| *S. cerevisiae* BY4741*Δhis3::HIS3-Pglk1-mTurquoise2-Tglk1 Δarg1:kanMX*$^{R}$ | *Sc*$^{CF50\ 3}$ | This work |
| *S. cerevisiae* BY4741*Δhis3::HIS3-Pglk1-mTurquoise2-Tglk1 Δarg1:kanMX*$^{R}$ | *Sc*$^{CF50\ 1\ Agg}$ | This work |
| *S. cerevisiae* BY4741*Δhis3::HIS3-Pglk1-mTurquoise2-Tglk1 Δarg1:kanMX*$^{R}$ | *Sc*$^{CF50\ 2\ Agg}$ | This work |
| *S. cerevisiae* BY4741*Δhis3::HIS3-Pglk1-mTurquoise2-Tglk1 Δarg1:kanMX*$^{R}$ | *Sc*$^{CF50\ 3\ Agg}$ | This work |
| *S. cerevisiae* BY4741*Δhis3::HIS3-Pglk1-mTurquoise2-Tglk1 Δarg1:kanMX*$^{R}$ | *Sc*$^{CF50\ 4\ Agg}$ | This work |
| *S. cerevisiae* BY4741*Δhis3::HIS3-Pglk1-mTurquoise2-Tglk1 Δarg1:kanMX*$^{R}$ | *Sc*$^{CF50\ 5\ Agg}$ | This work |
| *S. cerevisiae* BY4741*Δhis3::HIS3-Pglk1-mTurquoise2-Tglk1 Δarg1:kanMX*$^{R}$ | *Sc*$^{CF269\ 1}$ | This work |

| | | |
|---|---|---|
| *S. cerevisiae* BY4741*Δhis3::HIS3-Pglk1-mTurquoise2-Tglk1 Δarg1:kanMX*$^{R}$ | *Sc*$^{\text{CF50 2}}$ | This work |
| *S. cerevisiae* BY4741*Δhis3::HIS3-Pglk1-mTurquoise2-Tglk1 Δarg1:kanMX*$^{R}$ | *Sc*$^{\text{CF50 3}}$ | This work |
| *S. cerevisiae* BY4741*Δhis3::HIS3-Pglk1-mTurquoise2-Tglk1 Δarg1:kanMX*$^{R}$ | *Sc*$^{\text{CF50 4}}$ | This work |
| *S. cerevisiae* BY4741*Δhis3::HIS3-Pglk1-mTurquoise2-Tglk1 Δarg1:kanMX*$^{R}$ | *Sc*$^{\text{CF50 1 Agg}}$ | This work |
| *S. cerevisiae* BY4741*Δhis3::HIS3-Pglk1-mTurquoise2-Tglk1 Δarg1:kanMX*$^{R}$ | *Sc*$^{\text{CF50 2 Agg}}$ | This work |
| *S. cerevisiae* BY4741*Δhis3::HIS3-Pglk1-mTurquoise2-Tglk1 Δarg1:kanMX*$^{R}$ | *Sc*$^{\text{CF50 3 Agg}}$ | This work |
| *S. cerevisiae* BY4741*Δhis3::HIS3-Pglk1-mTurquoise2-Tglk1 Δarg1:kanMX*$^{R}$ | *Sc*$^{\text{CF50 4 Agg}}$ | This work |
| *S. cerevisiae* BY4741*Δhis3::HIS3-Pglk1-mTurquoise2-Tglk1 Δarg1:kanMX*$^{R}$ | *Sc*$^{\text{S 1}}$ | This work |
| *S. cerevisiae* BY4741*Δhis3::HIS3-Pglk1-mTurquoise2-Tglk1 Δarg1:kanMX*$^{R}$ | *Sc*$^{\text{S 2}}$ | This work |
| *S. cerevisiae* BY4741*Δhis3::HIS3-Pglk1-mTurquoise2-Tglk1 Δarg1:kanMX*$^{R}$ | *Sc*$^{\text{S 3}}$ | This work |
| *S. cerevisiae* BY4741*Δhis3::HIS3-Pglk1-mTurquoise2-Tglk1 Δarg1:kanMX*$^{R}$ | *Sc*$^{\text{S 4}}$ | This work |
| *S. cerevisiae* BY4741*Δhis3::HIS3-Pglk1-mTurquoise2-Tglk1 Δarg1:kanMX*$^{R}$ | *Sc*$^{\text{S 5}}$ | This work |
| *S. cerevisiae* BY4741*Δhis3::HIS3-Pglk1-mTurquoise2-Tglk1 Δarg1:kanMX*$^{R}$ | *Sc*$^{\text{S 6}}$ | This work |
| *S. cerevisiae* BY4741*Δhis3::HIS3-Pglk1-mTurquoise2-Tglk1 Δarg1:kanMX*$^{R}$ | *Sc*$^{\text{S 7}}$ | This work |
| *S. cerevisiae* BY4741*Δhis3::HIS3-Pglk1-mTurquoise2-Tglk1 Δarg1:kanMX*$^{R}$ | *Sc*$^{\text{S 8}}$ | This work |
| *S. cerevisiae* BY4741*Δhis3::HIS3-Pglk1-mTurquoise2-Tglk1 Δarg1:arg1* | *Sc*$^{\text{CF269 1 Arg1+}}$ | This work |
| *S. cerevisiae* BY4741*Δhis3::HIS3-Pglk1-mTurquoise2-Tglk1 Δarg1:arg1* | *Sc*$^{\text{CF269 2 Arg1+}}$ | This work |
| *S. cerevisiae* BY4741*Δade1:kanMX*$^{R}$ | | [54] |
| *S. cerevisiae* BY4741*Δade4:kanMX*$^{R}$ | | [54] |
| *S. cerevisiae* BY4741*Δade6:kanMX*$^{R}$ | | [54] |
| *S. cerevisiae* BY4741*Δade8:kanMX*$^{R}$ | | [54] |
| *S. cerevisiae* BY4741*Δarg4:kanMX*$^{R}$ | | [54] |
| *S. cerevisiae* BY4741*Δlys1:kanMX*$^{R}$ | | [54] |
| *S. cerevisiae* BY4741*Δlys4:kanMX*$^{R}$ | | [54] |
| *S. cerevisiae* BY4741*Δlys9:kanMX*$^{R}$ | | [54] |
| *S. cerevisiae* BY4741*Δser1:kanMX*$^{R}$ | | [54] |

| | | |
|---|---|---|
| *S. cerevisiae* BY4741*Δthr1:kanMX*$^R$ | | [54] |
| *S. cerevisiae* BY4741*Δtrp3:kanMX*$^R$ | | [54] |
| *S. cerevisiae* BY4741*Δtrp4:kanMX*$^R$ | | [54] |
| *S. cerevisiae* BY4741*Δhis3::HIS3-Pglk1-mTurquoise2-Tglk1 Δade1:kanMX*$^R$ | | This work |
| *S. cerevisiae* BY4741*Δhis3::HIS3-Pglk1-mTurquoise2-Tglk1 Δade4:kanMX*$^R$ | | This work |
| *S. cerevisiae* BY4741*Δhis3::HIS3-Pglk1-mTurquoise2-Tglk1 Δade6:kanMX*$^R$ | | This work |
| *S. cerevisiae* BY4741*Δhis3::HIS3-Pglk1-mTurquoise2-Tglk1 Δade8:kanMX*$^R$ | | This work |
| *S. cerevisiae* BY4741*Δhis3::HIS3-Pglk1-mTurquoise2-Tglk1 Δarg4:kanMX*$^R$ | | This work |
| *S. cerevisiae* BY4741*Δhis3::HIS3-Pglk1-mTurquoise2-Tglk1 Δlys1:kanMX*$^R$ | | This work |
| *S. cerevisiae* BY4741*Δhis3::HIS3-Pglk1-mTurquoise2-Tglk1 Δlys4:kanMX*$^R$ | | This work |
| *S. cerevisiae* BY4741*Δhis3::HIS3-Pglk1-mTurquoise2-Tglk1 Δlys9:kanMX*$^R$ | | This work |
| *S. cerevisiae* BY4741*Δhis3::HIS3-Pglk1-mTurquoise2-Tglk1 Δser1:kanMX*$^R$ | | This work |
| *S. cerevisiae* BY4741*Δhis3::HIS3-Pglk1-mTurquoise2-Tglk1 Δthr1:kanMX*$^R$ | | This work |
| *S. cerevisiae* BY4741*Δhis3::HIS3-Pglk1-mTurquoise2-Tglk1 Δtrp3:kanMX*$^R$ | | This work |
| *S. cerevisiae* BY4741*Δhis3::HIS3-Pglk1-mTurquoise2-Tglk1 Δtrp4:kanMX*$^R$ | | This work |

**Supplementary Table 6.** Plasmids used in this study

| **Plasmids** | **Addgene catalogue number** | **Reference** |
|---|---|---|
| pOB2 | | [65] |
| pNB1 | | [66] |
| pUA66 | | [67] |
| pSIJ8 | #68122 | [52] |
| pPL5071 | #60930 | [68] |
| pH3FS | #85780 | [69] |
| pBAD-lssmOrange | #37129 | [70] |
| pKD45 | | J. S. Parkinson, personal gift |
| pGS62 (pTrc99A:mNeonGreen) | | This work |
| pGS63 (pTrc99A:mCherry) | | This work |
| pGS64 (pTrc99A:mTurquoise2) | | This work |
| pGS65 (pTrc99A:lss-mOrange) | | This work |
| pGS66 (pUA66 $P_{hisJ}$ -86$_{A\text{->}T}$:GFP) | | This work |
| pGS67 (pUA66 $P_{hisJ}$ -87$_{A\text{->}T}$:GFP) | | This work |
| pGS68 (pUA66 $P_{hisJ}$ -78Δ:GFP) | | This work |
| pGS69 (pUA66 $P_{hisJ}$ -66Δ:GFP) | | This work |
| pGS70 (pUA66 $P_{hisJ}$ WT:GFP) | | This work |
| pGS5 (*HIS3-Pglk1-mTurquoise2-Tglk1)* | | [17] |
| pMFM073 (*HIS3-Pglk1-mNeonGreen-Tglk1)* | | [71] |

**Supplementary Table 7.** Primers used in this study

| **Primers** | **Sequence 5'->3'** | **Description** |
|---|---|---|
| GS_245 | cagaatttgcatgccgtgac | *ΔargR:kan$^R$* cassette amplification from the keio strain, from the evolved lines and for Sanger sequencing |
| GS_246 | ccttatgtattcattgtgtgaatgac | |
| GS_140 | catcctgactagtctttcaggc | *hisG:kan$^R$* cassette amplification from the keio strain |
| GS_177 | caaacttcgcgctgtattcc | |
| GS_281 | cctttcgtcttcacctcgagacggcacctacgacaagatg | Primers used to amplify the hisJ promoter to construct the plasmid pUA66:$P_{hisJ}$ :GFP reporters |
| GS_282 | tctccttcttaaatctagaggatccttaaaccagagagagcgatagcac | |
| pUA_fw | ggatcctctagatttaagaaggaga | Amplification of the backbone from the pUA:GFP plasmid |
| pUA_rv | ctcgaggtgaagacgaaagg | |
| GS_259 | aaatagagaagaacaagcaagatttttccctacccctattgggcatgccggtcgacggatctgatatcacc | *ecm21* KO cassette amplification for pH3FS |
| GS_260 | attcattcttcatcactcatcaaaggcactatttcgtcataacgcggaggatggtgtcgacaacccttaat | |
| GS_261 | gcattattctaatataacagttaggagaccaaaaagaaaaagaaatgtcagtcgacggatctgatatcacc | *gdh1* KO cassette amplification for pH3FS |
| GS_262 | agactatttaaaatacatcaccttggtcaaacatagcatcagagaccttgatggtgtcgacaacccttaat | |
| GS_332 | aagcatgccagtgttgaaatcagacaatttcgatccattggaagaagcttatggtgtcgacaacccttaat | *glt1* KO cassette amplification for pH3FS |
| GS_333 | tgactagctaattctttcaatagtttgtaatcacgttgaacgataccaccgtcgacggatctgatatcacc | |
| GS_363 | aatgacaagcgaaccagagtttcagcaggcttacgatgagatcgtttcttatggtgtcgacaacccttaat | *gdh3* KO cassette amplification for pH3FS |
| GS_364 | agcgcttacggctaaaaaacgtctccctggtcaagcattgcgtcagccacgtcgacggatctgatatcacc | |
| GS_367 | catacaaaacaaggatattaaattcacaacaataaaaagaataaagaatgatggtgtcgacaacccttaat | *gdh2* KO cassette amplification for pH3FS |
| GS_368 | attgaagctcaagcacttgcctccgcttctctttttaacccaccaataaagtcgacggatctgatatcacc | |
| GS_288 | cgatctggctgcaggacgtctggatgctgcgttacaagatgaagttgctggagcctgacatttatattcc | Amplification of the *neo-ccdB* cassette to replace the promoter region of *hisJ* |
| GS_289 | atggcgtcaatcttcttcgcttttaaggacgggattaacgcatccagcgggtcccgctcagaagaactc | |

#### Supplementary Data

**Supplementary Data 1. (separate file):** Original *E. coli* proteomics data presented in Extended Data Fig. 5

**Supplementary Data 2. (separate file):** Original *S. cerevisiae* proteomics data from the same co-cultures described in Extended Data Fig. 5

**Supplementary Data 3. (separate file):** Original *S. cerevisiae* proteomics data for Fig. 2i

**Supplementary Data 4. (separate file):** Complete mutation dataset obtained from *breseq* for *E. coli*

**Supplementary Data 5. (separate file):** Complete mutation dataset obtained from *breseq* for *S. cerevisiae*
